# CHD4 conceals aberrant CTCF-binding sites at TAD interiors by regulating chromatin accessibility in mESCs

**DOI:** 10.1101/2021.06.16.448663

**Authors:** Sungwook Han, Hosuk Lee, Andrew J. Lee, Seung-Kyoon Kim, Inkyung Jung, Gou Young Koh, Tae-Kyung Kim, Daeyoup Lee

**Affiliations:** Department of Biological Sciences, Korea Advanced Institute of Science and Technology, Daejeon, 34141, Republic of Korea; Center for Vascular Research, Institute for Basic Sciences, Daejeon, 34141, Republic of Korea; Department of Life Sciences, Pohang University of Science and Technology, Pohang 37673, Republic of Korea

**Keywords:** CHD4, ATP-dependent chromatin remodeler, mouse embryonic stem cells (mESCs), chromatin accessibility, CTCF, CTCF-binding sites, 3D chromatin organization, topologically associated domains (TADs), TAD borders, intrinsically disordered domains, B2 short interspersed nuclear element (SINE)

## Abstract

CTCF plays a critical role in the 3D chromatin organization by determining the TAD borders. Although CTCF primarily binds at the TAD borders, there also exist putative CTCF-binding sites within TADs, which are spread throughout the genome by retrotransposition. However, the detailed mechanism responsible for masking these putative CTCF-binding sites remains elusive. Here, we show that the ATP-dependent chromatin remodeler, CHD4, regulates chromatin accessibility to conceal aberrant CTCF-binding sites embedded in H3K9me3-enriched heterochromatic B2 SINEs in mouse embryonic stem cells (mESCs). Upon CHD4 depletion, these aberrant CTCF-binding sites become accessible, and aberrant CTCF recruitment occurs at the TAD interiors, resulting in disorganization of the local TADs. Furthermore, RNA-binding intrinsically disordered domains of CHD4 is required to prevent the aberrant CTCF bindings. Lastly, CHD4 is required for the repression of B2 SINE transcripts. These results highlight the CHD4-mediated mechanism that safeguards the appropriate CTCF bindings and associated TAD organizations in mESCs.

## Introduction

The mammalian genome is spatially organized into three-dimensional (3D) chromatin organizations, encompassing chromosome territories^1^, compartments (A and B compartments that are enriched for active/open chromatin and inactive/closed chromatin, respectively)^2,3^, topologically associated domains (TADs)^4,5^, contact/loop domains^6,7^, and insulated neighborhoods^8,9^, that allow for proper gene regulation. Previous studies found that the insulator protein, CCCTC-binding factor (CTCF), coexists with the cohesin complex in chromatin^10^ and localizes to the anchors/borders of chromatin loops^4–6,11^. Furthermore, removal of the cohesin loader, Nipbl^12^, loss of cohesin itself^7^, deletion of CTCF-binding sites^13–15^, or loss of CTCF itself^16^ have all been shown to interfere with the maintenance of 3D chromatin organizations, suggesting that these factors all play critical functions. Notably, the recent loss of function studies using single-cell analysis performed in mammalian cells reported two interesting observations^17,18^: the loss of CTCF resulted in the loss of TAD borders, resulting in the merging of two previously insulated TADs, whereas interactions within TAD were not significantly affected. On the other hand, the loss of cohesin subunit RAD21 caused significant disruptions in the interactions within the TAD, whereas TAD borders were not significantly affected. These findings demonstrated that 3D chromatin organizations are regulated by the combined action of CTCF and cohesin via distinct mechanisms: cohesin generates intra-TAD contacts, whereas CTCF prevents inter-TAD contacts by determining the TAD borders.

Importantly, in addition to the CTCF-binding sites (or motifs) that exhibit convergent orientation at the border of 3D domains^6^, there also exist putative CTCF-binding sites at the TAD interiors, which are spread throughout the genome by retrotransposition of B2 short interspersed nuclear element (SINE) retrotransposons^19,20^. Furthermore, from an evolutionary perspective, conserved CTCF-binding sites are enriched at the border of conserved 3D domains, while CTCF-binding sites that diverge between species (divergent CTCF-binding sites) drive local 3D structural changes at the domain interiors^21^. Thus, it is critical to mask the putative CTCF-binding sites that lie within the TADs because, similar to the divergent CTCF-binding sites, they could cause aberrant CTCF recruitment and disrupts the normal 3D chromatin organizations. Thus, although most studies focus on the function of CTCF at the TAD borders, preventing inappropriate CTCF bindings at the TAD interiors is equally essential for the maintenance of 3D chromatin organizations. However, the detailed mechanism responsible for masking these putative CTCF-binding sites remains poorly understood.

To address this issue, we focused on the chromodomain-helicase-DNA binding protein (CHD) family, especially CHD4, a well-known ATP-dependent chromatin remodeler. The CHD family is evolutionarily conserved^22^, and its members contribute to diverse cellular processes by assembling nucleosomes^23–26^. Since the majority of CTCF-binding sites are located at nucleosome-free regions^27^, it has been proposed that chromatin remodelers could govern CTCF recruitment by regulating nucleosome occupancy at certain CTCF-binding sites^28^. Furthermore, chromatin remodelers’ ability to regulate access to DNA through nucleosome positioning may contribute to obscure putative CTCF-binding sites.

Here, we show that CHD4 regulates chromatin accessibility to conceal aberrant CTCF-binding sites at the TAD interior, thereby preventing aberrant CTCF recruitments and securing the 3D chromatin organizations in mESCs. We use various NGS assays, including *in situ* Hi-C, MNase-seq, ATAC-seq, and H3/CTCF ChIP-seq, and perform temporal depletion/restoration of CHD4 to confirm the order of events; CHD4 initially assembles core histones to conceal aberrant CTCF-binding sites and by doing so prevents the aberrant CTCF bindings. We also identified that RNA-binding intrinsically disordered domains of CHD4 are required for preventing aberrant CTCF recruitments. Finally, we found that the CHD4-regulated aberrant CTCF-binding sites are embedded in H3K9me3-enriched heterochromatic B2 SINE retrotransposons, and CHD4 is required for the repression of B2 SINE transcripts. Together, our results demonstrate the detailed biological functions of CHD4 and elucidate the CHD4-modulated mechanism that secures the appropriate CTCF recruitments and TAD organizations in mESCs.

## Results

### CHD4 and CTCF are closely linked in mESCs

We first determined the expression of *Chd* family members using real-time quantitative polymerase chain reaction (RT-qPCR). The majority of the *Chd* expression levels were higher in mouse embryonic stem cells (mESCs) than in lineage-specified cells, such as MEFs or NIH3T3 cells (Supplementary Fig. 1A). Among the *Chd* family, *Chd4* was the most abundant in mESCs (Supplementary Fig. 1A), indicating that it could play an important role in these cells. Therefore, we focused on the biological function of CHD4 in mESCs.

To elucidate the function of CHD4, we first identified the CHD4-binding sites by performing cleavage under targets and release using nuclease (CUT&RUN)^29^ and also by analyzing the public chromatin immunoprecipitation sequencing (ChIP-seq) data against CHD4 (GSE64825, GSE61188, and GSE27844, see also Supplementary Table 1). Then, we analyzed genomic features within the CHD4-binding sites (CUT&RUN and ChIP-seq peaks). Notably, we found that CTCF motifs were highly enriched within CHD4-binding sites relative to CHD1-binding sites and randomly selected sites (Supplementary Fig. 1B, Supplementary Table 2). Interestingly, we also observed that CHD4 localizes to the border of TADs (Fig. 1a, Supplementary Fig. 1C) and anchor of insulated neighborhoods (Fig. 1b, Supplementary Fig. 1D), where CTCF is known to be localized (Fig. 1a, b) and play a boundary function^4–6,8,17,18^. Furthermore, 34% of the CTCF ChIP-seq peaks coincide with the CHD4 CUT&RUN peaks (Fig. 1c), suggesting that CHD4 and CTCF reside nearby. To further dissect this, we analyzed the CHD4 localization near the CTCF ChIP-seq peaks. As expected, we found that CHD4 resides near CTCF (Fig. 1d, e). Intriguingly, although CHD4-binding sites are enriched with CTCF motifs (Supplementary Fig. 1B, Supplementary Table 2), we observed the presence of CHD4 where CTCF is absent (Fig. 1c, e), indicating that CHD4 may conceal CTCF motifs (Fig. 1f). We also observed that CTCF localizes to exposed DNA sequences surrounded by well-positioned nucleosomes (Fig. 1d, e), consistent with the previous study^27^. Since CTCF is present at nucleosome-free regions flanking nucleosomes and CHD4 (Fig. 1d), this may suggest that CHD4 could regulate nucleosome occupancy near CTCF-binding sites, controlling interactions between CTCF and DNA sequence that harbor CTCF motifs. Taken together, these results strongly indicate that CHD4 and CTCF are closely linked in mESCs. Thus, we hypothesized that CHD4 might control the CTCF-mediated 3D chromatin architecture by regulating the chromatin accessibility near CTCF-binding sites (Fig. 1f).

**Fig. 1.**
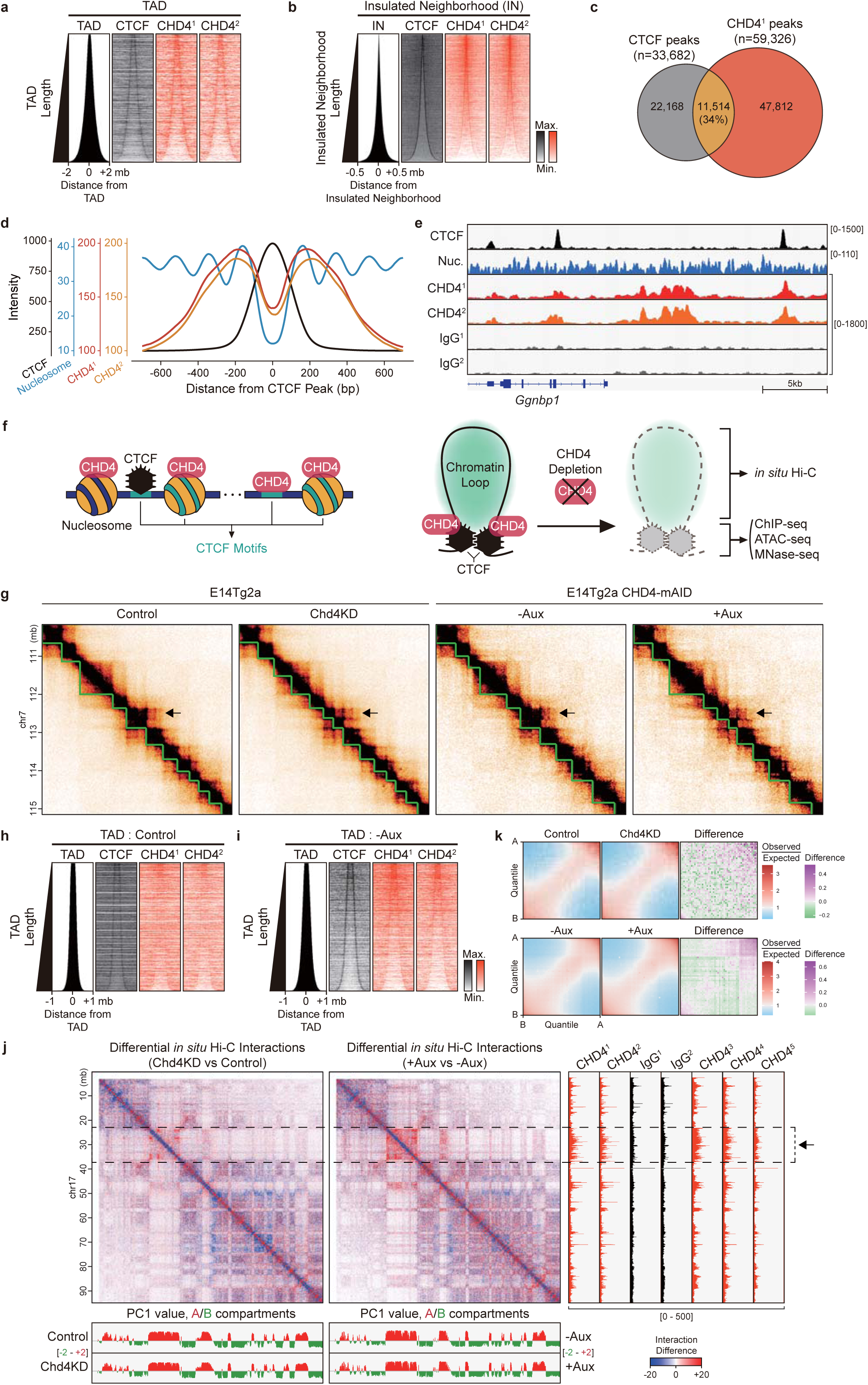
CHD4 localizes near CTCF and marginally organizes the global 3D chromatin architectures. **a, b** Heatmaps of CTCF and CHD4 aligned at 2,153 TADs^4^ (**a**) and 23,726 insulated neighborhoods^8^ (**b**). All heatmaps were sorted in ascending order by the length of the TAD or insulated neighborhood. CHD4^1^ (in-house generated antibody) and CHD4^2^ (Abcam antibody) denote CUT&RUN data using different antibodies against CHD4. See also Supplementary Fig. 1C, D. **c** Venn diagrams representing the overlap of CTCF and CHD4^1^ peaks (binding sites). **d** Line plots showing average enrichments of CTCF, nucleosomes, and CHD4 at CTCF peaks. **e** Examples representing the position/enrichment of CTCF, nucleosomes (Nuc.), CHD4^1,2^ and IgG^1,2^ (which are relevant to CHD4^1,2^). **f** Model in which CTCF motifs are enriched within CHD4-binding sites, and CTCF is localized at nucleosome-free regions flanking nucleosomes and CHD4 (left). Model showing that both CHD4 and CTCF are localized at the boundary regions of chromatin loops (right). After CHD4 depletion, we conducted the listed experiments to detect changes in the 3D chromatin architecture (*in situ* Hi-C), CTCF enrichment (ChIP-seq), and chromatin accessibility (ATAC-seq and MNase-seq). **g** Examples of Hi-C interactions (heatmaps) and TADs (green lines). Black arrows indicate the region where 3D chromatin architectures are severely disrupted upon CHD4 depletion. **h, i** Heatmaps of CTCF and CHD4^1,2^ aligned at 5,452 TADs in Control cells (**h**) and 4,525 TADs in auxin-untreated (-Aux) cells (**i**). All heatmaps were sorted in ascending order by the length of the TADs. **j** Examples of differential Hi-C interactions and CHD4 localization at chromosome 17. Distribution of PC1 (first principal components, equivalent to the first eigenvectors) values across the chromosomes and A (red) / B (green) compartments are shown (bottom). Black arrow and dotted box indicate the CHD4-enriched regions where 3D chromatin architectures are disrupted upon CHD4 depletion. **k** *cis* PC1 values in 40 kb genomic bins are ranked into 50 quantiles (A and B compartments), and pairwise Hi-C enrichment (compartment strength) was calculated between each of the 50 quantiles. The difference in compartment strength upon CHD4 depletion was calculated (right).

### CHD4 depletion marginally affects genome-wide 3D chromatin architectures

To test our hypothesis, we first designed siRNAs against *EGFP* (Control) and *Chd4* (Chd4KD). We confirmed the successful knockdown of *Chd4* in mESCs using RT-qPCR (Supplementary Fig. 1E) and immunoblot (Supplementary Fig. 1F). Furthermore, we employed an auxin-inducible degron (AID) system^30^ to degrade CHD4 in mESCs (CHD4-mAID) (Supplementary Fig. 1G-L). We confirmed that CHD4 was efficiently degraded after 24 hours of auxin treatment using immunoblot (Supplementary Fig. 1J, 4B).

To investigate whether CHD4 depletion disturbed the 3D chromatin architecture, we performed *in situ* Hi-C upon *Chd4* knockdown (Control and Chd4KD cells) and CHD4 depletion (auxin untreated and treated cells, termed as -Aux and +Aux cells, respectively). We generated a total of ∼2.1 billion valid Hi-C interactions (Supplementary Table 3) and comprehensively analyzed *in situ* Hi-C data using HiC-Pro^31^ and GENOVA^32^. The Hi-C experiments were performed in two biological replicates and showed high reproducibility according to the Pearson correlation coefficient (Supplementary Fig. 1M). Furthermore, we applied reciprocal insulation scores to identify TADs (Fig. 1g) by using CaTCH^33^. Consistent with previous results (Fig. 1a, b), we observed that CHD4 and CTCF both localize at the border of Control cells’ TADs (Fig. 1h) and –Aux cells’ TADs (Fig. 1i). Notably, we found changes in TAD positions (Fig. 1g, arrow) and changes in Hi-C interactions at the CHD4-enriched regions (Fig. 1j, dotted box with arrow) upon CHD4 loss. However, we did not detect major changes in relative contact probability (RCP) (Supplementary Fig. 1N) and changes in compartment domains (Fig. 1j, Supplementary Fig. 1O). In addition, we observed that only ∼2.3 or ∼2.5% of compartment domains (in 100-kb bins) are switched to the opposite one (from A to B or from B to A compartment) upon CHD4 depletion (Supplementary Fig. 1P). Interestingly, we detected a slight increase in AA compartmentalization strength upon CHD4 loss (Fig. 1k). Despite changes in Hi-C interactions, this largely-unaffected compartmentalization is consistent with previous studies^7,16^. Together, we concluded that CHD4 depletion marginally affects global 3D chromatin architectures.

### CHD4 maintains the local 3D chromatin architecture by preventing aberrant CTCF recruitment at TAD interiors

To investigate the role of CHD4 in the local 3D chromatin architecture, we performed aggregate TAD analysis (ATA) upon CHD4 depletion at four TADs obtained from each cell (Control, Chd4KD, -Aux, and +Aux cells). Notably, we detected decreased Hi-C interactions within Control and –Aux cells’ TADs in a genome-wide manner upon CHD4 depletion (Fig. 2a, b, arrow). In contrast, Hi-C interactions within Chd4KD and +Aux cells’ TADs were largely unaltered by CHD4 loss (Fig. 2a, b). These results suggest that CHD4 maintains the local interactions within the original TADs of wild-type mESCs.

**Fig. 2.**
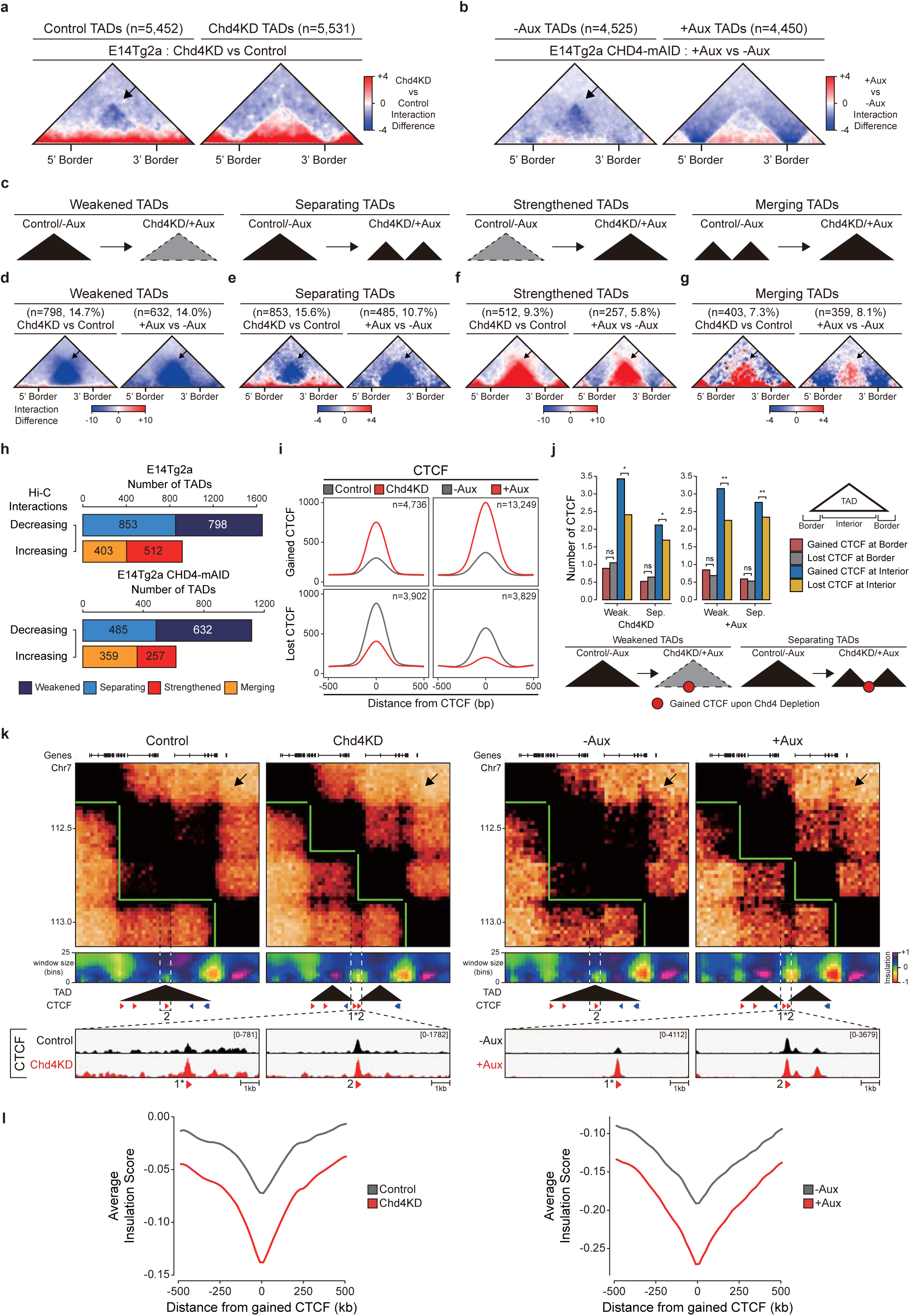
CHD4 maintains the local 3D chromatin architectures by ensuring proper CTCF recruitment. **a, b** Aggregate TAD analysis (ATA) comparing Hi-C interactions of Chd4KD and Control cells at the 5,452 Control TADs (left) and 5,531 Chd4KD TADs (right) (**a**) and +Aux and –Aux cells at the 4,525 –Aux TADs (left) and 4,450 +Aux TADs (right) (**b**). **c** Schematic diagrams showing the four types of modified TADs upon CHD4 depletion. **d-g** ATA results at weakened (**d**), separating (**e**), strengthened (**f**), and merging (**g**) TADs. **h** Stacked bar graphs showing the number of decreasing and increasing TADs upon *Chd4* knockdown (top) and CHD4 depletion (bottom). **i** Line plots showing average enrichments of gained CTCF (top) and lost CTCF (bottom) upon CHD4 depletion. **j** Bar graphs showing the average number of gained and lost CTCF at the TAD border and interior. The average number of CTCF was normalized with respect to the total number of relevant CTCF types. *P*-values were derived using Wilcoxon signed rank test (\**P* < 1×10^−10^; \*\**P* < 1×10^−50^; ns, not significant). **k** Example of separating TADs upon CHD4 depletion. Hi-C interactions (heatmaps), TADs (green lines and black triangles), insulation heatmaps, CTCF ChIP-seq peaks (red and blue triangles indicate + and – directionalities of CTCF motifs), and genes are shown in Control, Chd4KD, -Aux, and +Aux cells. Black arrows indicate the region where 3D chromatin architectures are severely disrupted upon CHD4 depletion. The zoom-in CTCF ChIP-seq data represent the gained CTCF (1*) which resides at a newly-established border of separating TADs (dotted boxes) upon CHD4 depletion. See also Supplementary Fig. 2B, C. **l** Line plots showing average insulation scores at the gained CTCF upon CHD4 depletion.

To address this further, by comparing the changes in Hi-C interactions, we identified weakened, strengthened TADs, while by comparing the changes in TAD positions, we identified separating, merging TADs; thus, we identified four types of differential TADs upon CHD4 depletion (Fig. 2c-g). Weakened TADs were defined as original TADs of wild-type mESCs (Control or –Aux cells), in which their Hi-C interactions were decreased upon CHD4 loss. In contrast, the strengthened TADs were defined as TADs of CHD4-depleted cells (Chd4KD or +Aux cells), which their Hi-C interactions were increased upon CHD4 loss. When considering the changes in TAD positions, separating TADs were defined as original TADs of wild-type mESCs (Control or –Aux cells) that were split into two or more smaller TADs upon CHD4 loss. Reversely, merging TADs were defined as TADs of CHD4-depleted cells (Chd4KD or +Aux cells) that contain two or more original TADs of wild-type mESCs (Control or –Aux cells). According to the ATA results, we observed that Hi-C interactions were genome-widely decreased at the weakened TADs (Fig. 2d) and separating TADs (Fig. 2e), while increased at the strengthened TADs (Fig. 2f) and merging TADs (Fig. 2g) upon CHD4 loss. Regarding the TAD positions, we found that decreasing TADs (weakened and separating TADs) share more TADs compared to increasing TADs (strengthened and merging TADs) (Supplementary Fig. 2A), indicating that weakened and separating TADs are closely associated. Interestingly, by comparing the number of four differential TADs upon CHD4 loss, we found that decreasing TADs occurs more frequently than increasing TADs (Fig. 2h). These more frequent occurrences of decreasing TADs reflect our previous observation of genome-wide decreases in Hi-C interactions within the original TADs of wild-type mESCs (Fig. 2a, b, arrow). Thus, we focused on decreasing TADs (weakened and separating TADs).

To elucidate how CHD4 maintains the local interactions within TADs, we performed CTCF ChIP-seq upon CHD4 loss. By comparing the CTCF intensity, we defined gained and lost CTCF upon CHD4 depletion (Fig. 2i). Next, we divided decreasing TADs (weakened and separating TADs) into borders and interiors (Fig. 2j, top right). We observed that both weakened and separating TADs exhibit similar distribution patterns of gained/lost CTCF at the border/interior of TADs (Fig. 2j), suggesting that these two decreasing TADs may be fundamentally the same or maybe formed similarly upon CHD4 loss. At TAD borders, we did not detect significant differences in the distribution of gained and lost CTCF (Fig. 2j). In contrast, we found that gained CTCF was significantly more distributed at TAD interiors compared to lost CTCF (Fig. 2j). These results indicate that gained CTCF may be responsible for decreasing interactions within weakened and separating TADs upon CHD4 loss.

To further support this, we comprehensively analyzed Hi-C interactions, TAD positions, insulations scores, and CTCF ChIP-seq upon CHD4 loss at specific regions. Consistent with the previous genome-wide ATA results (Fig. 2d, e), we found a notable decrease in Hi-C interactions within the weakened and separating TADs (Fig. 2k, Supplementary Fig. 2B, C, black arrow). Importantly, we observed that gained CTCF (marked with *) greatly reduced the local insulation, thereby generating a novel TAD border (separating TAD, Fig. 2k, Supplementary Fig. 2B) and/or markedly decreasing the Hi-C interactions within TADs (weakened TAD, Supplementary Fig. 2B, C). In general, we observed the same types of decreasing TAD upon CHD4 loss (Chd4KD and +Aux cells), but in a particular case, we observed weakened TAD in Chd4KD cells and separating TAD in +Aux cells at the same genomic loci (Supplementary Fig. 2B). This observation supports the idea that these two decreasing TADs are closely associated; the gained CTCF causes both types of decreasing TADs upon CHD4 depletion. When considering the splitting feature of separating TADs (Fig. 2c), one can speculate the reduced TAD sizes and an elevated number of chromatin loops (peaks) coupled with gained CTCF upon CHD4 loss. As expected, we observed reduced TAD sizes (Supplementary Fig. 2D) and an elevated number of total chromatin loops (peaks) and cell-type-specific peaks (Supplementary Fig. 2E, F) upon CHD4 depletion (Supplementary Fig. 2G, H). Importantly, at the gained CTCF sites, we detected that insulation scores were globally reduced upon CHD4 depletion (Fig. 2l), indicating that CHD4 depletion-triggered gained CTCF is responsible for directly generating TAD borders in a genome-wide manner. Furthermore, our aggregate peak analysis (APA) results showed the increasing Hi-C interactions at the gained CTCF-associated peaks upon CHD4 loss (Supplementary Fig. 2I), supporting the direct relationship between gained CTCF and changes in Hi-C interactions. Collectively, our results strongly imply that decreasing TADs (weakened and separating TADs) are directly associated with gained CTCF upon CHD4 loss. Therefore, these findings suggest that CHD4 regulates the local 3D chromatin architecture by preventing aberrant CTCF recruitment at TAD interiors.

### CHD4 conceals aberrant CTCF-binding sites by regulating chromatin accessibility at heterochromatic regions

To investigate how CHD4 prevents aberrant CTCF binding, we analyzed the genome-wide changes in CTCF upon CHD4 depletion at CHD4 CUT&RUN peaks (or CHD4-binding sites) (Fig. 3a, b, Supplementary Fig. 3A, B). We classified the CHD4 CUT&RUN peaks into those that coincide with CTCF ChIP-seq peaks of wild-type mESCs (Control or –Aux cells) and those that do not coincide. We did not detect any major changes in CTCF upon CHD4 depletion at CHD4 peaks that coincide with CTCF peaks (Fig. 3a, b, Supplementary Fig. 3A, B). In contrast, we detected aberrantly gained CTCF upon CHD4 depletion at CHD4 peaks that do not coincide with CTCF peaks (Fig. 3a, b, Supplementary Fig. 3A, B, arrow). Since CHD4 CUT&RUN peaks are highly enriched with CTCF motifs (Supplementary Fig. 1B, Supplementary Table 2), these results indicate that CHD4 conceals putative CTCF motifs (Fig. 1f, left). The observed changes in CTCF binding were not derived from any change in *CTCF* expression (Supplementary Fig. 3C, D).

**Fig. 3.**
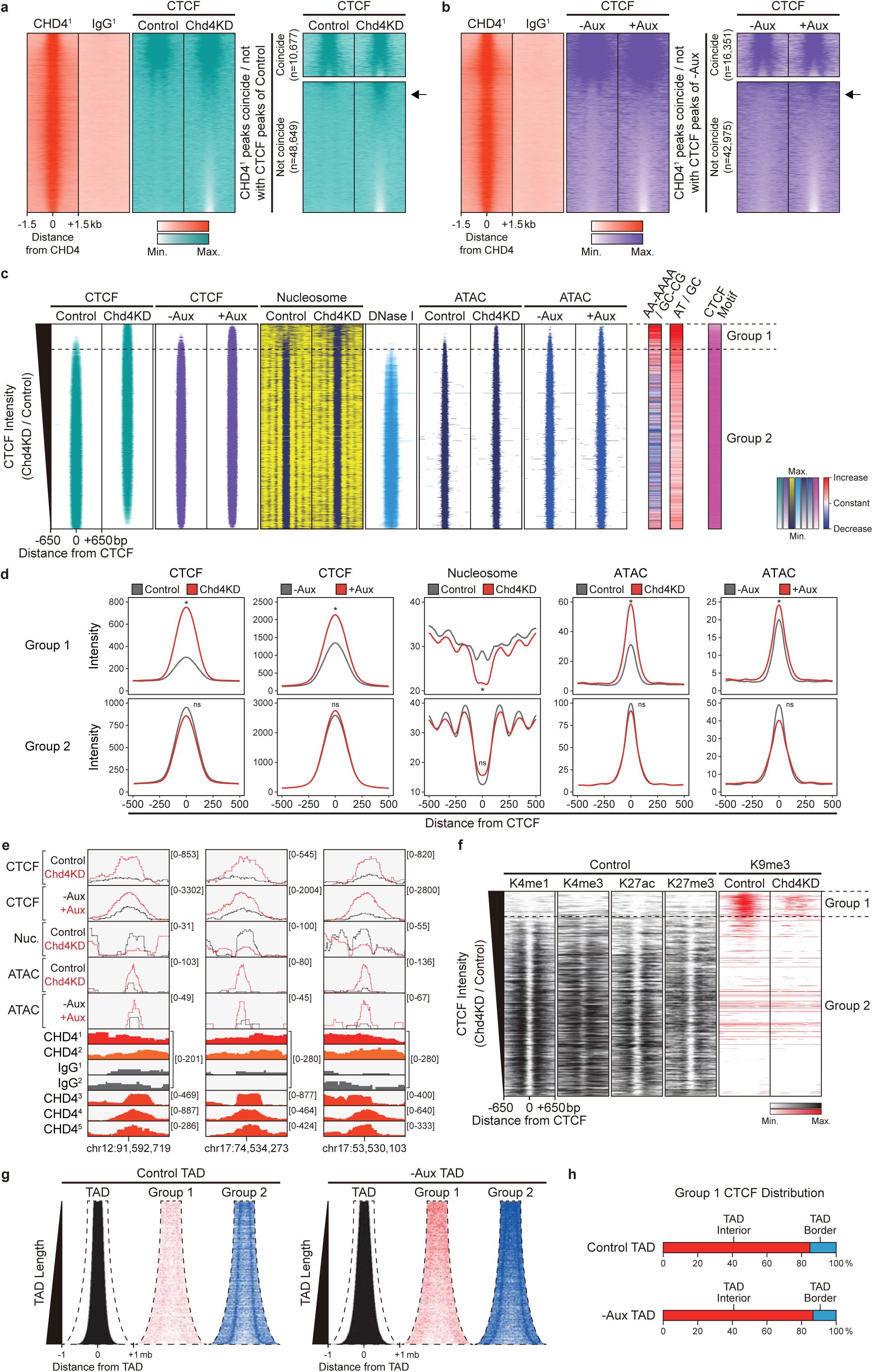
CHD4 conceals aberrant CTCF-binding sites at TAD interiors by regulating chromatin accessibility. **a, b** Heatmaps representing CHD4^1^, IgG^1^ CUT&RUN, CTCF ChIP-seq of Control and Chd4KD cells (**a**), and CTCF ChIP-seq of –Aux and +Aux cells (**b**) at CHD4 peaks (left), CHD4 peaks that coincide (top right), and not coincide with CTCF peaks (bottom right). **c** Heatmaps displaying CTCF, nucleosome, DNase I hypersensitive sites (DNase I), ATAC-seq signals (ATAC), the ratio of nucleosome-repelling sequences (AA-AAAA) or DNA bases (AT) to nucleosome-preferring sequences (GC-CG) or DNA bases (GC), and CTCF motifs at all CTCF peaks in Control and Chd4KD cells as indicated on top. See also Supplementary Fig. 3F. **d** Line plots showing average enrichments of CTCF, nucleosome, and ATAC at Group 1 and Group 2 CTCF-binding sites. *P*-values were derived using Wilcoxon signed rank test (\**P* < 1×10^−100^; ns, not significant). **e** Examples of Group 1 sites, including representative results for CTCF, nucleosomes (Nuc.), ATAC, CHD4, IgG. **f** Heatmaps displaying various histone H3 modifications in Control cells (black) and heterochromatin marker, H3K9me3, in Control and Chd4KD cells (red) as indicated on top. See also Supplementary Fig. 3H. **c, f** All heatmaps were aligned at 39,800 CTCF peaks (rows) and sorted in descending order by the relative CTCF intensity (Chd4KD/Control). 4,736 Group 1 (11.9%) represents CTCF-binding sites at which more CTCF were recruited in Chd4KD cells compared to Control cells (1.75-fold), while 35,064 Group 2 (88.1%) represents the remainder of the CTCF-binding sites, at which CTCF binding was unchanged or reduced in Chd4KD cells. Heatmaps that are not labeled as Control or Chd4KD represent wild-type mESCs. **g** Heatmaps representing TAD positions (black), Group 1 (red), and Group 2 (blue) CTCF-binding sites at Control TADs (left) and –Aux TADs (right). All heatmaps were sorted in ascending order by the length of the TADs. **h** Percentage bar graph showing the distribution of Group 1 CTCF-binding sites at the interior and border of Control TADs (top) and –Aux TADs (bottom).

To elucidate how CHD4 conceals putative CTCF motifs, we comprehensively analyzed CTCF ChIP-seq data obtained from Control and Chd4KD cells. We first determined the CTCF-binding sites (or ChIP-seq peaks) in Control and Chd4KD cells. We then merged the CTCF-binding sites (n=39,800) and sorted them in descending order by their relative CTCF intensity (Chd4KD/Control) (Fig. 3c, f, Supplementary Fig. 3E, H-J). According to this relative CTCF intensity, we defined two groups of CTCF-binding sites: Group 1 (n=4,736, 11.9%) represents CTCF-binding sites at which more CTCF were recruited in Chd4KD cells compared to Control cells, while Group 2 (n=35,064, 88.1%) represents the remainder of the CTCF-binding sites, at which CTCF binding was unchanged or reduced in Chd4KD cells. Consistent with the previous results (Fig. 2i, k, 3a, b, Supplementary Fig. 2B, C, E, 3A, B), we observed that CHD4 depletion-triggered gain and loss of CTCF binding occurred at the Group 1 and Group 2 CTCF-binding sites, respectively (Fig. 3c). Furthermore, we found that RAD21 is aberrantly gained along with CTCF at the Group 1 sites upon CHD4 depletion (Supplementary Fig. 3E, left). This supports our previous results that aberrantly gained CTCF upon CHD4 depletion is directly involved in forming new chromatin loops (Supplementary Fig. 2I), thereby disrupting local 3D chromatin organizations.

Previously, we observed that CTCF binds to exposed DNA sequences surrounded by well-positioned nucleosomes (Fig. 1d)^27^, and CHD4 is also localized near these sites (Fig. 1d). In addition, we detected the aberrantly gained CTCF upon CHD4 depletion at the CHD4 CUT&RUN peaks (Fig. 3a, b, Supplementary Fig. 3A, B), which harbor CTCF motifs. Considering these results, we hypothesized that CHD4 conceals putative CTCF motifs by regulating chromatin accessibility at specific DNA sequences that embed CTCF motifs (Fig. 1f, left). To examine whether this is the case, we analyzed MNase-seq (chromatin digestion with micrococcal nuclease combined with sequencing), DNase-seq (DNase I hypersensitive sites sequencing), and ATAC-seq (assay for transposase-accessible chromatin using sequencing) data and aligned them with the same order as previously defined relative CTCF intensity (Group 1 and 2 sites). In Control cells, CTCF mainly localized to Group 2 sites (Fig. 3c), which were characterized by an accessible chromatin region with depleted nucleosomes, enriched DNase I hypersensitive sites, and enriched ATAC-seq signals (Fig. 3c). In accordance with this, Control cells exhibited far less binding of CTCF at Group 1 sites (Fig. 3c), which were characterized by an inaccessible chromatin region with enriched nucleosomes, depleted DNase I hypersensitive sites, and depleted ATAC-seq signals (Fig. 3c). Importantly, we detected a significant decrease in nucleosomes and, reversely, a significant increase in ATAC-seq signals upon CHD4 depletion at Group 1 sites (Fig. 3c-e), indicating that CHD4 depletion caused the Group 1 sites to change from inaccessible chromatin regions to accessible chromatin regions. In contrast, we did not detect any significant changes in chromatin accessibility at Group 2 sites (Fig. 3c, d).

To examine why nucleosomes were depleted specifically at Group 1 sites, we analyzed the features of these DNA sequences. Notably, we found that nucleosome-repelling sequences (AA-AAAA) and DNA bases (AT)^34–37^ were more abundant within Group 1 sites compared to Group 2 sites (Fig. 3c). Moreover, CTCF motifs were also present within Group 1 sites (Fig. 3c), even though these sites were not typically targeted by CTCF in Control cells (Fig. 3c). Consistently, when we performed the same analysis using CTCF ChIP-seq peaks of –Aux and +Aux cells, we observed similar changes in CTCF, RAD21, and ATAC-seq signals upon CHD4 depletion at Group 1* sites (Supplementary Fig. 3F, G). The Group 1* sites also exhibited similar patterns (depleted DNase I hypersensitive sites, enriched nucleosome-repelling sequences, and enriched CTCF motifs) as Group 1 sites (Fig. 3c). Taken together, our results suggest that CHD4 maintains the inaccessible/closed chromatin state and conceals CTCF motifs at Group 1 sites by assembling nucleosomes, thereby preventing aberrant CTCF recruitment. Upon CHD4 depletion, however, nucleosomes are no longer assembled by CHD4 and instead tend to be removed from Group 1 sites due to the highly enriched nucleosome-repelling sequences, resulting in the open/accessible chromatin states, exposure of CTCF motifs, and the mislocalization of CTCF. Thus, our data support the idea that the CHD4-mediated regulation of chromatin accessibility controls the appropriate recruitment of CTCF in mESCs.

We next determined the epigenetic features of the Group 1 sites using ChIP-seq data against various factors, including histone modifications. Consistent with enriched nucleosomes (Fig. 3c), we observed enriched histone H3.1/2 at Group 1 sites in wild-type mESCs (Supplementary Fig. 3H). Remarkably, among the various histone modifications, H3K9me3 (Fig. 3f, Supplementary Fig. 3H) and the H3K9 methyltransferases, Suv39h1 and Suv39h2, but not Setdb1 (Supplementary Fig. 3H), were the only marks enriched at Group 1 sites in wild-type mESCs. As expected, given that the chromatin structure changed from the closed (enriched nucleosomes) to open chromatin state (depleted nucleosomes) at Group 1 sites upon CHD4 depletion, H3K9me3 was also diminished at these regions in Chd4KD cells (Fig. 3f) due to the absence of core histones, substrates for H3K9 methyltransferases. Moreover, this reduction was not related to any change in the expression of genes encoding H3K9 methyltransferases (Supplementary Table 4). Furthermore, we observed that only CHD4 (Supplementary Fig. 3E) and ChAHP complexes (Supplementary Fig. 3I) were abundant at Group 1 sites. In contrast, most of the analyzed chromatin remodelers (Supplementary Fig. 3E), mediators, histone variants, histone modifications (Supplementary Fig. 3H), NuRD complexes, architectural proteins, and transcription factors (Supplementary Fig. 3J) were sparse at the Group 1 sites. Together, these results indicate that CHD4 assembles core histones to conceal putative CTCF motifs at heterochromatic regions.

Lastly, we analyzed the positions of the Group 1 and Group 2 CTCF-binding sites with respect to the TADs. We observed that Group 2 sites were mostly localized at the TAD borders (Fig. 3g). In contrast, we found that Group 1 sites (equivalent to gained CTCF in Fig. 2i) were not located at the TAD borders (Fig. 3g). Instead, they were mainly located at the interior regions of Control cells’ TADs (84.8%) and –Aux cells’ TADs (86.7%) (Fig. 3h), consistent with our previous results (Fig. 2j, k, Supplementary Fig. 2B, C). Furthermore, when considering our reduced insulation scores (Fig. 2l) and APA results (Supplementary Fig. 2I), these data collectively imply that CHD4 depletion-triggered gained CTCF (Group 1 sites) is directly associated with the disruption of local TAD organizations by forming a novel TAD border and/or chromatin loops at TAD interiors.

### CHD4 initially assembles core histones to conceal aberrant CTCF-binding sites and thereby prevents the aberrant CTCF bindings

Previously, we found that CHD4 depletion increased both chromatin accessibility (decreased nucleosome levels and increased ATAC-seq signals) and CTCF occupancy at Group 1 sites (Fig. 3, Supplementary Fig. 3). Since CHD4 is a chromatin remodeler, we speculated that CHD4 depletion would cause changes in chromatin accessibility first, leading to the exposure of CTCF motifs, then CTCF is newly recruited to the Group 1 sites, resulting in aberrant CTCF bindings (Fig. 3, Supplementary Fig. 3, see also Fig. 4a). To confirm this, we carried out two types of temporal aspect experiments to determine the order of events (Fig. 4a, h). We first performed temporal depletion of CHD4 by varying the RNAi (siRNA against *Chd4*) treatment time (Fig. 4b). As our expectation, we observed a gradual decrease in *Chd4* expression levels as we increased the RNAi treatment time (Fig. 4c). At these time points, we performed ChIP-seq against CTCF and histone H3. Consistent with previous results (Fig. 3, Supplementary Fig. 3), we did not detect any major changes in CTCF and H3 levels at Group 2 sites upon gradual depletion of CHD4 (Fig. 4d). Notably, we observed the gradual increase in CTCF levels (gained CTCF) and gradual decrease in H3 levels (becoming more accessible chromatin states) at Group 1 sites upon temporal depletion of CHD4 (Fig. 4d). To assess these accurately, we calculated the average intensity of CTCF and H3 at Group 1 sites at each time point (Fig. 4e, f). Importantly, as our original speculation, we found that H3 levels decrease first between 6-12hr (hours) of CHD4 depletion (Fig. 4f, red arrow), and then CTCF levels increase later on between 12-48hr of CHD4 depletion (Fig. 4e, red arrow). These results were also confirmed when we examined the individual loci of Group 1 sites (Fig. 4g, Supplementary Fig. 4A), indicating that upon CHD4 depletion, changes in chromatin accessibility (closed to open chromatin state) occur first, followed by aberrantly gained CTCF bindings.

**Fig. 4.**
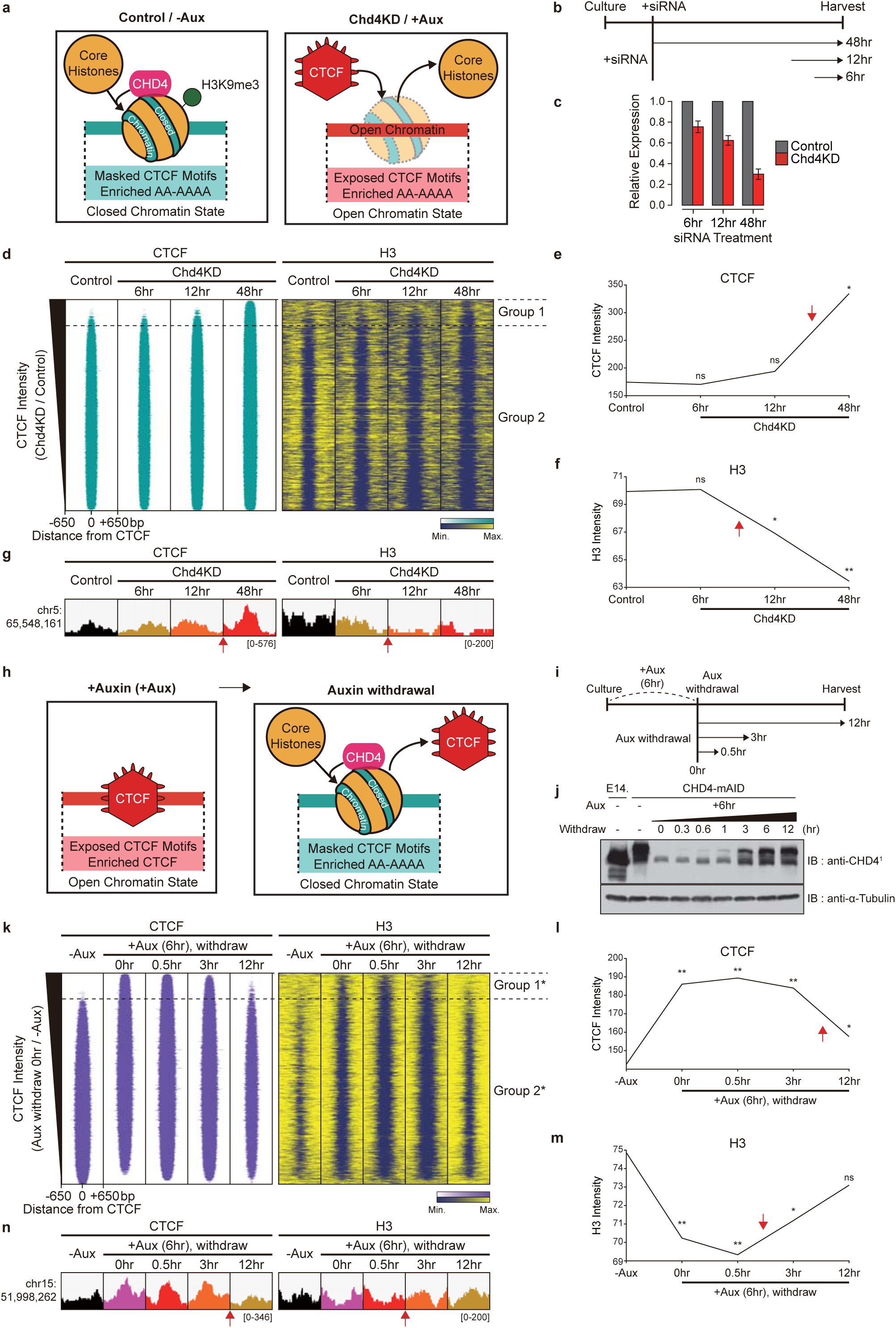
CHD4 first assembles core histones and thereby prevents the aberrant CTCF binding at the Group 1 sites. **a** Model illustrating the role of CHD4 at the Group 1 sites based on our results in Fig. 3 and Supplementary Fig. 3. **b** Schematic representation of temporal aspect experiments with various siRNA (against *Chd4*) treatment time. **c** RT-qPCR analysis of *Chd4* expression after various siRNA treatment times. Error bars denote the standard deviation obtained from three biological replicates. The expression levels were normalized with respect to that of β-actin. **d** Heatmaps displaying CTCF and H3 ChIP-seq of various siRNA treatment times at all CTCF peaks in Control and Chd4KD cells as indicated on top. All heatmaps were aligned and sorted in the same way as described in Fig. 3c. See also Fig. 3c (siRNA against *Chd4* treated for 48 hours). **e** Line plots showing average enrichments of CTCF at Group 1 sites. *P*-values were derived using Wilcoxon signed rank test (\**P* < 1×10^−100^; ns, not significant). **f** Line plots showing average enrichments of H3 at Group 1 sites. *P*-values were derived using Wilcoxon signed rank test (\**P* < 1×10^−3^; \*\**P* < 1×10^−15^; ns, not significant). **e, f** *P*-values were calculated by comparing the Control and each of Chd4KD time points. **g** Examples of Group 1 sites, including representative results for CTCF and H3 across a time course of siRNA treatment. See also Supplementary Fig. 4A. **h** Model representing the aberrant CTCF binding upon CHD4 depletion (left) and our prediction after CHD4 restoration via auxin withdrawal (right). **i** Schematic representation of auxin withdrawal experiments with various time points. **j** Immunoblots of CHD4 after auxin treatment (6 hours) followed by various auxin withdrawal times in CHD4-mAID cells and E14Tg2a wild-type (E14.). α-Tubulin was detected as a loading control. **k** Heatmaps displaying CTCF and H3 ChIP-seq of –Aux and +Aux followed by various auxin withdrawal times at all CTCF peaks in -Aux and +Aux (auxin withdraw 0 hr) cells as indicated on top. All heatmaps were aligned at 59,327 CTCF peaks (rows) and sorted in descending order by the relative CTCF intensity (auxin withdraw 0 hr/-Aux). 7,225 Group 1* (12.2%) represents CTCF-binding sites at which more CTCF were recruited in +Aux (auxin withdraw 0 hr) compared to –Aux cells (1.5-fold), while 52,102 Group 2* (87.8%) represents the remainder of the CTCF-binding sites. See also Supplementary Fig. 3F (auxin was treated for 24 hours) and 4C. **l** Line plots showing average enrichments of CTCF at Group 1* sites. *P*-values were derived using Wilcoxon signed rank test (\**P* < 1×10^−100^; \*\**P* < 1×10^−300^; ns, not significant). **m** Line plots showing average enrichments of H3 at Group 1* sites. *P*-values were derived using Wilcoxon signed rank test (\**P* < 1×10^−30^; \*\**P* < 1×10^−50^; ns, not significant). **l, m** *P*-values were calculated by comparing the -Aux and each of auxin withdrawal time points. **n** Examples of Group 1* sites, including representative results for CTCF and H3 across a time course of auxin withdrawal. See also Supplementary Fig. 4D. **e-g, l-n** Red arrows indicate the time points when intensity (CTCF or H3) changes most rapidly after siRNA treatment (**e-g**) or auxin withdrawal (**l-n**).

To validate our findings, we performed temporal restoration of CHD4 levels (Fig. 4h). We first confirmed that CHD4 is completely degraded after 6hr of auxin treatment in CHD4-mAID cells (Supplementary Fig. 4B). Then, we removed the auxin by changing the cell culture media and found that CHD4 protein is gradually restored during 12hr of auxin withdrawal (Fig. 4i, j). To determine whether core histones assemble first or CTCF eviction occurs first at Group 1 sites upon CHD4 restoration, we varied the auxin withdrawal time (Fig. 4i) and performed ChIP-seq against CTCF and histone H3. Consistent with previous results (Fig. 3, 4d, Supplementary Fig. 3), we found that CHD4 depletion, which is equivalent to 6hr of auxin treatment without auxin withdrawal (‘+Aux (6hr), withdraw 0hr’), caused the decreased H3 levels and increased CTCF levels at Group 1 sites (Supplementary Fig. 4C). Notably, we observed the gradual decrease in CTCF levels (gained CTCF) and gradual increase in H3 levels (becoming more accessible chromatin states) at Group 1 sites upon temporal restoration of CHD4 (Supplementary Fig. 4C). When we performed the same analysis using CTCF ChIP-seq peaks of –Aux and +Aux cells (‘+Aux (6hr), withdraw 0hr’), we observed the same rescuing patterns of CTCF and H3 levels (that is, the gradual decrease in CTCF levels and gradual increase in H3 levels) upon temporal restoration of CHD4 at Group 1* sites (Fig. 4k), as consistent at Group 1 sites (Supplementary Fig. 4C). To quantitatively assess these, we calculated the average intensity of CTCF and H3 at Group 1* sites at each time point (Fig. 4l, m). Importantly, we found that H3 levels increase first between 0.5-3hr of auxin withdrawal (Fig. 4m, red arrow), and then CTCF levels decrease later on between 3-12hr of auxin withdrawal (Fig. 4l, red arrow). Furthermore, these results were also confirmed when we examined the individual loci of Group 1* sites (Fig. 4n, Supplementary Fig. 4D), indicating that CHD4 restoration first rescues chromatin accessibility (open to closed chromatin state) then rescues CTCF bindings. Collectively, we determined the order of events and concluded that CHD4 directly regulates chromatin accessibility at the Group1 sites to conceal putative CTCF motifs and thereby prevents aberrant CTCF bindings.

### RNA-binding intrinsically disordered domain of CHD4 is required to prevent aberrant CTCF bindings

We next questioned why CHD4 depletion majorly affects the chromatin accessibility at the Group 1 sites while exhibiting modest changes at the Group 2 sites, although CHD4 is bound at both Group 1 and 2 sites (Supplementary Fig. 3E). The most plausible explanation for the modest changes at Group 2 sites upon CHD4 depletion would be the presence of various factors, such as chromatin remodelers (Supplementary Fig. 3E), mediators, histone variants, histone modifications (Supplementary Fig. 3H), NuRD complexes, architectural proteins, and transcription factors (Supplementary Fig. 3J). Thus, these factors, particularly chromatin remodelers (CHD1 and BRG1), may compensate for the loss of CHD4 at Group 2 sites. In contrast, at Group 1 sites, these other factors are sparse while CHD4, ChAHP complex and SUV39H1/2 are abundant. Together, these facts may explain why CHD4 depletion primarily affects the chromatin states of putative CTCF motifs at Group 1 sites.

Alternatively, differential chromatin states at Group 1 and 2 sites may explain these differential effects upon CHD4 depletion. As described previously, Group 1 sites represent heterochromatic regions, while Group 2 sites represent relatively euchromatic regions (Fig. 3c, f, Supplementary Fig. 3H) where RNAs are abundant. Since CHD4 is an RNA-binding protein^38,39^, we hypothesized that RNAs might inhibit the catalytic activity of CHD4 (allowing that CHD4 functions majorly at RNA-depleted heterochromatic Group 1 sites in wild-type mESCs), which may explain why CHD4 depletion affects primarily at Group 1 sites. We first confirmed that CHD4 antibodies could be used in immunoprecipitation (Supplementary Fig. 5A), performed RIP (RNA immunoprecipitation), and found that CHD4 bound with the *Oct4* and *Nanog* RNAs (Fig. 5a), which are very abundant in mESCs^40^. Furthermore, we also confirmed the RNA binding ability of purified CHD4 (Supplementary Fig. 5B) using RNA EMSAs (electrophoretic mobility shift assays) (Fig. 5b, Supplementary Fig. 5C). Together, these results demonstrated that CHD4 binds to RNAs, consistent with previous reports^38,39^. To determine the influence of RNA binding on the activity of CHD4, we performed *in vitro* nucleosome-sliding assays with various concentrations of RNA. Interestingly, we observed that the nucleosome-sliding activity of CHD4 decreased as the RNA concentration increased (Fig. 5c, Supplementary Fig. 5D). Collectively, our data suggest that RNAs bind to CHD4 and inhibit its catalytic activity, which may explain (along with other possibilities mentioned above) why CHD4 plays a modest role at RNA-abundant euchromatic Group 2 sites.

**Fig. 5.**
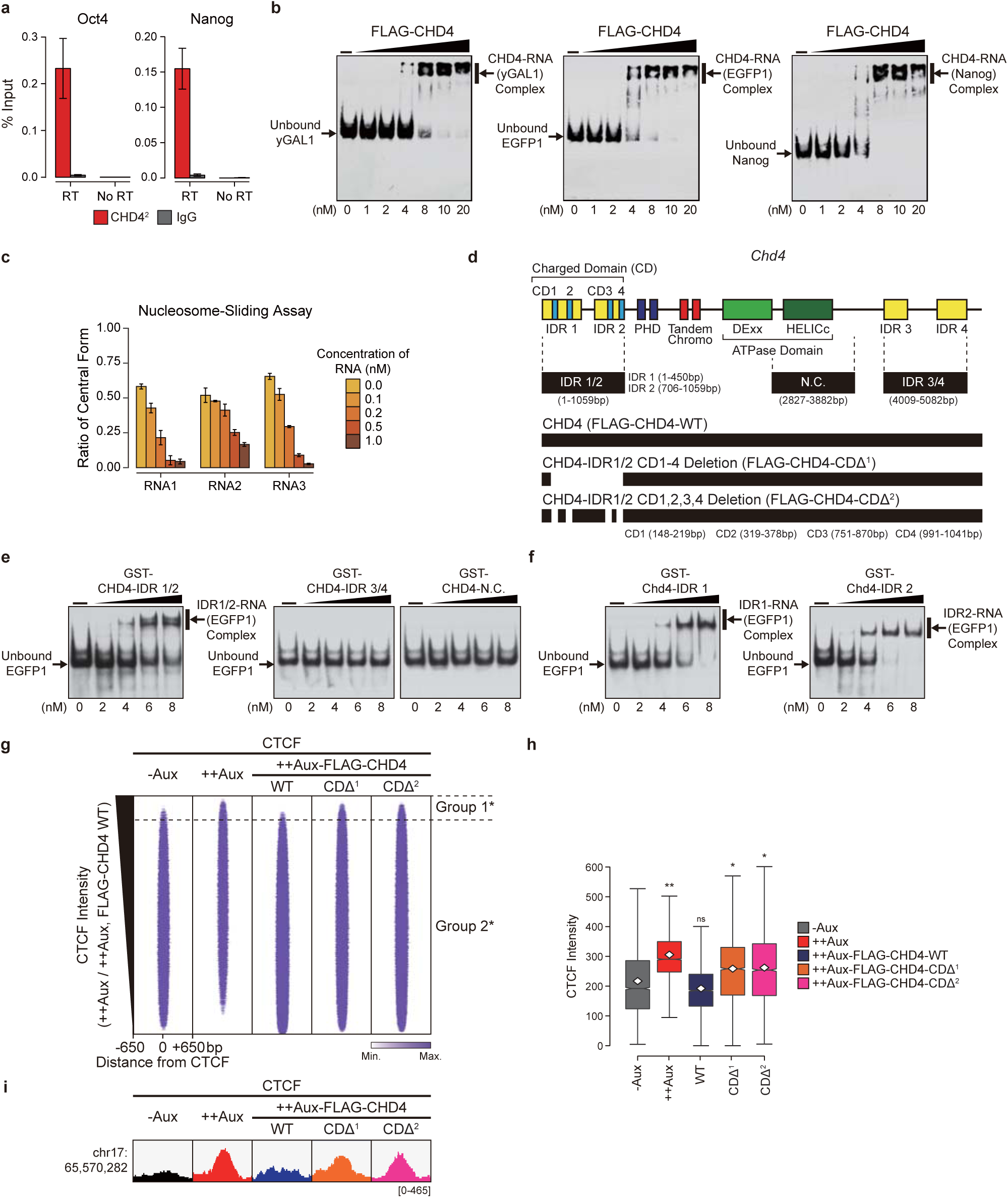
RNA binding ability of CHD4 is required to prevent aberrant CTCF binding at the Group 1 sites. **a** RNA immunoprecipitation (RIP) assay of Chd4. RIP was carried out with the indicated antibodies. All signals were normalized with respect to the input signal. Error bars denote the standard deviation obtained from three biological replicates. **b** Electrophoretic mobility shift assays (EMSAs) for the *yGAL1* (left), *EGFP1* (middle), and *Nanog* (right) RNAs were performed with purified FLAG-CHD4. See also Supplementary Fig. 5C. **c** Bar graphs showing the ratio of the central form to total nucleosomes (central and lateral forms) according to the results of nucleosome-sliding assays (Supplementary Fig. 5D). RNA1, 2, and 3 denote *yGAL1*, *EGFP1*, and *EGFP2*, respectively. Error bars denote the standard deviation obtained from three biological replicates. **d** Schematic representation of various domains within *Chd4* genes. Specific positions of domains that we used are indicated. **e, f** EMSAs for the *EGFP1* RNAs were performed with purified GST-CHD4-IDR 1/2 (**e**, left), GST-CHD4-IDR 3/4 (**e**, middle), GST-CHD4-N.C. (**e**, right), GST-CHD4-IDR 1 (**f**, left), and GST-CHD4-IDR 2 (**f**, right). **b, e, f** ‘Unbound RNA’ indicates free RNA, and ‘CHD4 (or IDR)-RNA Complex’ indicates the binding of the purified protein to each RNA species. **g** Heatmaps displaying CTCF ChIP-seq of –Aux, ++Aux, and ++Aux followed by various ectopic transfection of FLAG-CHD4 constructs at all CTCF peaks in ++Aux and ++Aux-FLAG-CHD4-WT cells as indicated on top. All heatmaps were aligned at 57,631 CTCF peaks (rows) and sorted in descending order by the relative CTCF intensity (++Aux /++Aux-FLAG-CHD4-WT). 4,326 Group 1* (7.5%) represents CTCF-binding sites at which more CTCF were recruited in ++Aux compared to ++Aux-FLAG-CHD4-WT cells (1.5-fold), while 53,305 Group 2* (92.5%) represents the remainder of the CTCF-binding sites. See also Supplementary Fig. 5E-G. **h** Box plots showing the enrichments of CTCF at Group 1* sites. *P*-values were calculated by comparing the -Aux and each of the other samples. The horizontal line and the white rhombus in the box denote the median and mean, respectively. *P*-values were derived using Wilcoxon signed rank test (\**P* < 1×10^−100^; \*\**P* < 1×10^−200^; ns, not significant). **I** Examples of Group 1* sites, including representative results for CTCF. See also Supplementary Fig. 5H.

To investigate the importance of RNA-binding ability of CHD4 in the regulation of CTCF bindings, we first aim to identify the specific RNA-binding domains of CHD4. Previously, it is known that many RNA-binding proteins contain intrinsically disordered regions (IDR), and the IDRs can bind to RNAs^41,42^. Regarding these, we used MobiDB^43,44^ to determine the RNA-binding IDR of CHD4. The MobiDB predicted four IDRs (IDR 1-4) within *Chd4* (Fig. 5d). We first purified three types of proteins, each containing IDR 1/2, IDR 3/4, and negative control regions (N.C.), and then performed RNA EMSAs. We found that only IDR 1/2 (1-1059bp of *Chd4*) exhibits the RNA binding ability (Fig. 5e). Notably, we found that both IDR 1 (1-450bp) and IDR 2 (706-1059bp) binds to RNAs (Fig. 5f). Thus, we specifically identified the RNA binding domains of CHD4. To exclude target-off effects, we narrowed down the regions within IDRs. Since protein-RNA interaction interfaces are known to be formed by clusters of positively charged residues that are scattered on protein surfaces^45,46^, we defined four positive charged residues (charged domain, CD 1-4) within IDR 1 and IDR 2 (Fig. 5d). We next created DNA constructs of *Chd4* mutants that cannot bind to RNAs by deleting all regions containing four charged domains (CD) of IDR 1 and IDR 2 (Fig. 5d, CDΔ^1^), and we also deleted all of individual CD (Fig. 5d, CDΔ^2^) to minimize the target-off effects. To perform rescue experiments, we initially transfected ectopic DNA constructs of FLAG-tagged CHD4 full-length (FLAG-CHD4-WT) and RNA binding defective CHD4 (FLAG-CHD4-CDΔ^1^ or -CDΔ^2^), and then treat auxin to deplete endogenous CHD4 proteins (Supplementary Fig. 5E, left, ‘+Auxin 24hr (+Aux)’). Interestingly, although FLAG-CHD4-CDΔ^1,2^ was successfully overexpressed (Supplementary Fig. 5F, left, see lane 6 and 7), we failed to overexpress FLAG-CHD4-WT (Supplementary Fig. 5F, left, see lane 3). We found that this failure was due to the presence of endogenous CHD4-mAID proteins, probably forming dimers with ectopic FLAG-CHD4-WT proteins and degrade together (in case of FLAG-CHD4-CDΔ^1,2^, it fails to form dimers with endogenous CHD4-mAID proteins, resulting in the stable expression). To resolve this, we treated auxin before transfection (Supplementary Fig. 5E, right, ‘++Auxin 48hr (++Aux)’), leading to degradation of endogenous CHD4-mAID proteins prior to the FLAG-CHD4-WT expressions. By doing so, we could observe the stable expression of FLAG-CHD4 WT proteins (Supplementary Fig. 5F, right, see lane 10). To elucidate the role of RNA-binding CD of CHD4 in the regulation of CTCF bindings, we performed rescue experiments by transfecting FLAG-CHD4-WT/-CDΔ^1^/-CDΔ^2^ to express equally (Supplementary Fig. 5G) after complete depletion of endogenous CHD4 proteins and then performed CTCF ChIP-seq. Consistent with our previous results (Fig. 3, 4), we observed aberrantly gained CTCF at Group 1* sites upon CHD4 depletion (++Aux) (Fig. 5g). Importantly, we found that ectopic transfection of FLAG-CHD4-WT (full-length CHD4) rescued the CTCF levels at Group 1* sites (Fig. 5g-i, Supplementary Fig. 5H), indicating that CHD4 is directly associated with preventing aberrant CTCF bindings at these sites. However, strikingly, we observed that CTCF levels were not entirely restored in both of *Chd4* mutants (FLAG-CHD4-CDΔ^1,2^) that cannot bind to RNAs (Fig. 5g-i, Supplementary Fig. 5H). Thus, we revealed that RNA-binding charged domain (CD) within intrinsically disordered domain (IDR) of CHD4 is required to prevent aberrant CTCF bindings in mESCs.

### CHD4 is required for the repression of B2 SINEs in mESCs

Previously, we defined the epigenetic features of H3K9me3-enriched heterochromatic aberrant CTCF-binding sites (Group 1) and determined the role of CHD4 at these sites. To elucidate the genomic features of Group 1 sites, we next examined the distribution of genomic regions by performing genome annotation. Notably, we found that SINEs (particularly B2 SINEs), which are well-known retrotransposons^47^, were enriched specifically at Group 1 sites (Fig. 6a); this is consistent with previous reports that H3K9me3 suppresses SINEs^48,49^, which harbor CTCF motifs^19,20^. Since CHD4 depletion caused the loss of nucleosomes/core histones (Fig. 3, 4, Supplementary Fig. 3, 4), resulting in the reduction of H3K9me3 levels (Fig. 3f) at Group 1 sites, we next investigated whether this affects the transcription of B2 SINE by performing mRNA-seq, total RNA-seq, and GRO (global run-on)-seq. Surprisingly, we observed that all of the signals obtained from the three assays were elevated upon CHD4 depletion at Group 1 sites (Fig. 6b, Supplementary Fig. 6A), where B2 SINEs are enriched (Fig. 6a). Furthermore, our GRO-seq analysis revealed over 2-fold increase in the number of *de novo* SINE transcripts upon CHD4 depletion (Supplementary Fig. 6B). Notably, our total RNA-seq analysis showed that the expression of B2 SINEs, localized at the Group 1 sites, were up-regulated upon CHD4 depletion (Supplementary Fig. 6C), strongly implying that CHD4 directly represses B2 SINE transcripts at Group 1 sites. As B2 SINE is mainly suppressed by H3K9me3 via Suv39h1 and Suv39h2 in mESCs^48,49^, our results indicate that B2 SINEs within Group 1 sites are repressed by enriched H3K9me3 and its methyltransferases in Control cells, but become de-repressed upon CHD4 depletion due to the reduction of a repressive mark, H3K9me3.

**Fig. 6.**
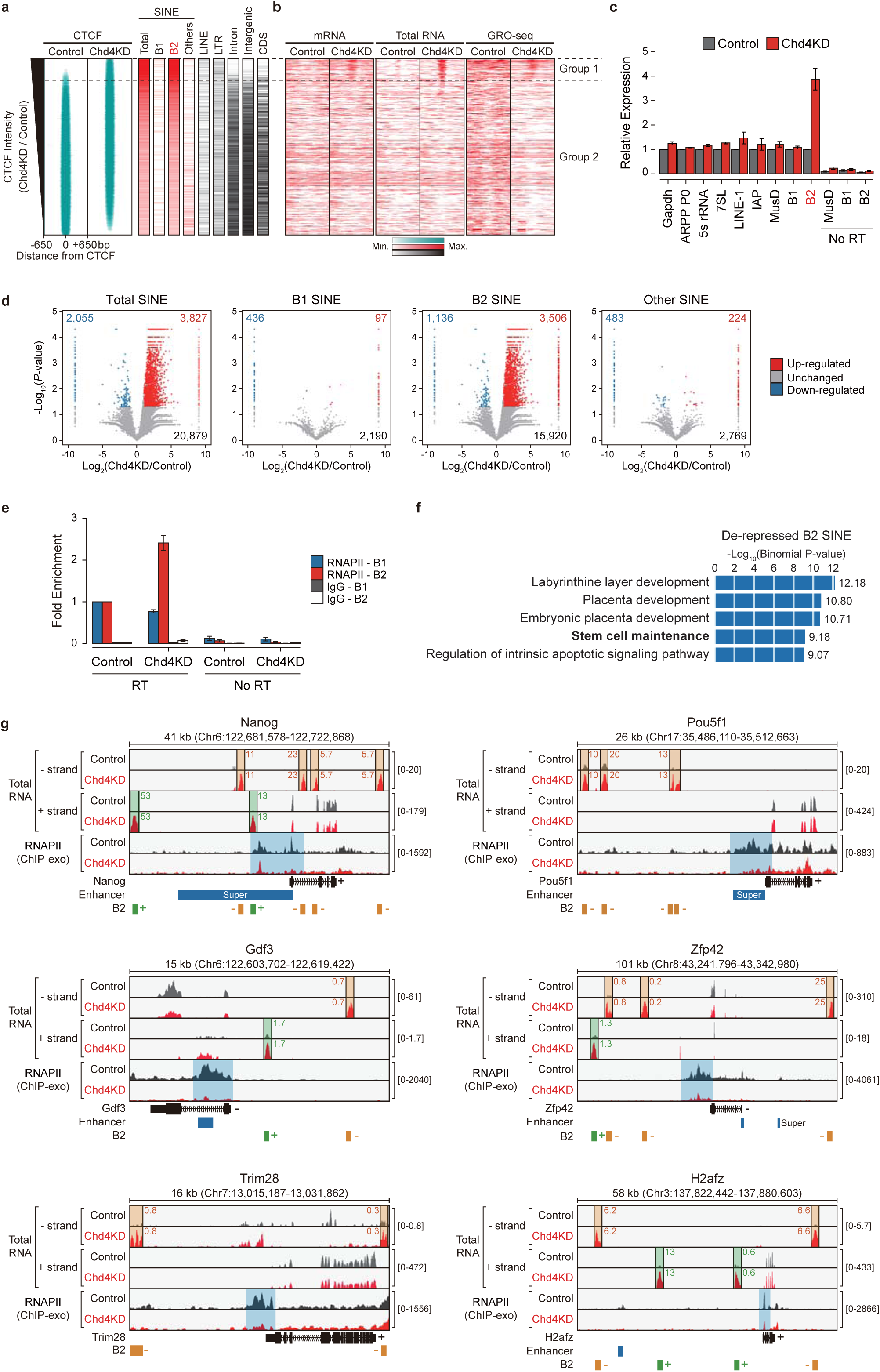
CHD4 represses B2 SINEs in mESCs. **a** Heatmaps displaying the genomic contents of the mouse genome. CDS includes the exon, promoter, transcription start site (TSS), and transcription termination site (TTS). **b** Heatmaps representing mRNA-seq, total RNA-seq, and GRO-seq data in Control and Chd4KD cells. **a, b** All heatmaps were aligned and sorted in the same way as described in Fig. 3c. See also Fig. 3 and Supplementary Fig. 3. **c** RT-qPCR analysis of retrotransposons in mESCs. Gapdh and ARPP are transcribed by RNAPII. The 5s rRNA, 7SL, and retrotransposons are mainly transcribed by RNAPIII. The obtained expression levels were normalized with respect to that of the 28S rRNA, which is transcribed by RNA Pol I. Error bars denote the standard deviation obtained from three biological replicates. **d** Volcano plots showing the differential expression of SINEs (Total, B1, B2, and other SINE), as measured using total RNA-seq. Only SINEs localized at intergenic regions were analyzed. Blue and red dots indicate the expressions of SINEs that are significantly decreased and increased, respectively, upon *Chd4* knockdown (*P*-value ≤ 0.05 and log_2_ (Chd4KD/Control) ≤ −1 or ≥ +1). **e** RNA immunoprecipitation (RIP) assay of B1 and B2 SINEs bound by RNAPII. Blue, red, gray, and white boxes indicate RNAPII (8WG16)-bound B1 SINE, RNAPII (8WG16)-bound B2 SINE, IgG-bound B1 SINE, and IgG-bound B2 SINE, respectively. Immunoprecipitated SINEs were normalized with respect to the levels of B1 or B2 SINEs bound by RNAPII in Control cells. Error bars denote the standard deviation obtained from three biological replicates. **f** Results of GREAT (Genomic Regions Enrichment of Annotations Tool) analysis, which was used to identify a class of genes that are located near de-repressed B2 SINEs (see Method). **g** Examples of de-repressed B2 SINEs (- and + strand total RNA-seq) and disrupted localization of RNAPII (ChIP-exo) at various stem cell maintenance-related genes. Orange, green, and blue boxes highlight changes in total RNA-seq (- strand), total RNA-seq (+ strand), and RNAPII (ChIP-exo), respectively. Genes, enhancers, and B2 SINE loci and their strand directions are marked at the bottom. See also Supplementary Fig. 6F.

To confirm this, we determined the expression levels of various retrotransposons, including LINEs, ERVs, and SINEs, using RT-qPCR. Consistently, the expression level of B2 SINE, a major mouse SINE, was dramatically up-regulated in Chd4KD cells, whereas the other retrotransposons (e.g., LINE-1 and IAP) were not significantly altered (Fig. 6c). Although B2 SINEs transcripts are known to be transcribed by RNA polymerase III, there are some B2 SINEs located within introns, which may be affected by RNA polymerase II (RNAPII). To exclude this possibility, we treated the RNAPII inhibitor α-amanitin and then determined the expression of B2 SINEs in mESCs. Consistent with previous results, we found that B2 SINEs were specifically up-regulated by *Chd4*-knockdown even when RNAPII was inhibited (Supplementary Fig. 6D). These results were also confirmed by our total RNA-seq analysis on SINEs that are localized at the intergenic regions of the mouse genome; we found that CHD4 depletion specifically up-regulated B2 SINE while not affecting B1 and other SINEs (Fig. 6d). Thus, these data indicate that CHD4 represses global B2 SINE transcripts in mESCs.

### CHD4 depletion-triggered de-repressed B2 SINE may hinder the RNA polymerase II recruitment at the transcription start sites of pluripotent genes

We next investigated the role of de-repressed B2 SINE upon CHD4 depletion in mESCs. As up-regulated B2 SINE RNA has been reported to directly bind RNAPII and inhibit transcription^50,51^, we examined whether the CHD4 depletion-triggered de-repressed B2 SINE exhibit the same features. We first performed RIP (RNA immunoprecipitation) and confirmed that de-repressed B2 SINE transcripts have a binding preference for RNAPII in Chd4KD cells (Fig. 6e). We then used GREAT (Genomic Regions Enrichment of Annotations Tool) analysis^52^ to identify the set of genes located in the vicinity of de-repressed B2 SINEs. Interestingly, our results revealed that genes that are important for stem cell maintenance (pluripotent genes) were located near de-repressed B2 SINEs (Fig. 6f), as well as Group 1 CTCF-binding sites (Supplementary Fig. 6E). To investigate whether de-repressed B2 SINE transcripts hinder the recruitment of RNAPII to the TSSs (transcription start sites) of genes involved in stem cell maintenance, we analyzed our total RNA-seq and public RNAPII ChIP-exo data (GSE64825, see also Supplementary Table 1). For detailed analysis, we separated the total RNA-seq data into those for the −/+ strands and confirmed the presence of de-repressed B2 SINEs in Chd4KD cells (Fig. 6g). Notably, Chd4KD cells exhibited a significant reduction of RNAPII recruitment at TSSs when abundant de-repressed B2 SINEs were located nearby (Fig. 6g), while no reduction of RNAPII recruitment was observed at TSSs that lacked a nearby de-repressed B2 SINE (Supplementary Fig. 6F). The observed changes in RNAPII recruitment were not derived from any change in the expression levels of the RNAPII subunits (Supplementary Fig. 6G). To determine whether this reduction of RNAPII recruitment at TSS could dysregulate the expression levels of stem cell maintenance-related genes, we analyzed total RNA-seq and mRNA-seq. Notably, we observed that genes that possess abundant de-repressed B2 SINEs, resulting in reduced RNAPII recruitment at their TSS (Fig. 6g), were down-regulated (Supplementary Fig. 6H, red) compared to the genes that lacked a nearby de-repressed B2 SINE (Supplementary Fig. 6H, gray). Taken together, our data indicate that *Chd4* knockdown induces de-repressed B2 SINE, which prevents the proper recruitment of RNAPII to the TSSs of stem cell maintenance genes (e.g., *Nanog* and *Oct4*), resulting in down-regulation of these genes. Thus, CHD4 is required to repress B2 SINEs, which may hinder the recruitment of RNAPII at TSS of pluripotent genes.

## Discussion

Here, we explore the changes in 3D chromatin architecture of mESCs upon CHD4 depletion using *in situ* Hi-C. We find that CHD4 critically contributes to the maintenance of TADs (Hi-C interactions within TADs and TAD positions) by preventing aberrant CTCF bindings at the TAD interior. Most importantly, we discover that CHD4 conceals aberrant CTCF-binding sites (Group 1 sites) by regulating chromatin accessibility, thereby maintaining these sites as inaccessible, H3K9me3-enriched heterochromatic regions. Thus, upon CHD4 depletion, the aberrant CTCF-binding sites become accessible, exposing the putative CTCF motifs, and aberrant CTCF recruitment occurs, resulting in Weakened/Separating TADs. We also show that these aberrant CTCF-binding sites are embedded in B2 SINEs, which are normally repressed by CHD4 in wild-type mESCs. Upon CHD4 depletion, B2 SINEs are de-repressed (due to the reduction of B2 SINE-repressive mark, H3K9me3) and interact with RNAPII, disrupting the RNAPII recruitment to the TSS of stem cell maintenance-related genes, resulting in dysregulation of relevant gene expressions. Collectively, our results elucidate a novel mechanism that CHD4 regulates chromatin accessibility at aberrant CTCF-binding sites, thereby safeguarding the appropriate CTCF bindings and maintaining the local TAD organizations in mESCs (Fig. 7).

**Fig. 7.**
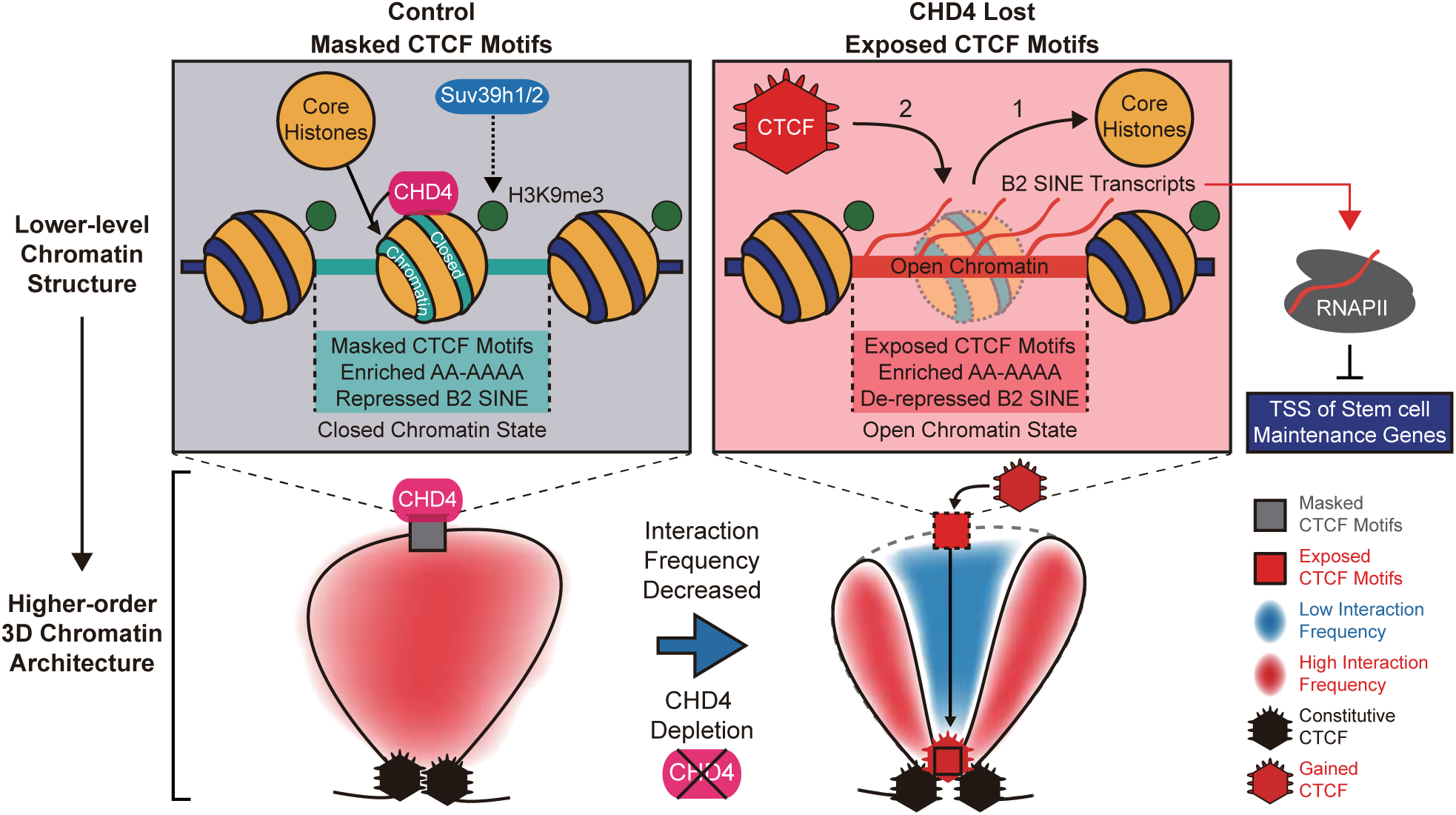
Proposed model illustrating the function of CHD4; CHD4 regulates the lower-level chromatin structures (top), thereby maintains the higher-order 3D chromatin architectures (bottom).

Since the genome is packaged into 3D chromatin organizations and most of the biological processes, including transcription, occur in this context, it is important to investigate the 3D organization of the genome. Previous studies characterized the 3D chromatin organization in terms of compartments^2,3^, TADs^4,5^, contact/loop domains^6,7^, and insulated neighborhoods^8,9^. Recent reports have mainly addressed the importance of CTCF and cohesin in 3D genome topology by observing the disruption of 3D chromatin structures after CRISPR/Cas9 system-mediated deletion/orientation change of CTCF-binding sites^13–15^, deletion of cohesin^7,17,18^, and CTCF^16–18^. These studies mainly focused on the properties and formation of the 3D chromatin architecture and the key factors (e.g., CTCF and cohesin) that compose its structures. However, intriguing questions remained, such as (1) How are TADs established or maintained? (2) How are aberrant CTCF binding and associated Hi-C interactions at TAD interiors prevented? (3) How is CTCF recruitment at the TAD borders regulated? (4) Does chromatin remodelers involve in CTCF recruitments? (5) What is the role of chromatin remodelers in 3D chromatin organizations? We believe that our present work provides clues to these questions by elucidating a CHD4-mediated mechanism that conceals aberrant CTCF-binding sites at the TAD interiors by regulating chromatin accessibility, thereby preventing inappropriate CTCF bindings and safeguarding both proper CTCF recruitment and 3D chromatin architecture in mESCs.

It is interesting to set our results in the context of evolutionary aspects. A previous report compared Hi-C data from four mammals and identified two types of CTCF sites: (1) conserved CTCF sites, which are conserved across species and are enriched at strong TAD borders; and (2) divergent CTCF sites, which lack conservation across species, are newly evolved, and localize at TAD interiors^21^. The authors also demonstrated that the evolutionary dynamics of intra-TAD interactions, which reflect the actions of divergent CTCF sites, could play a critical role in enhancer-promoter interactions within TADs^21^. In support of this, a recent review suggested that the 3D chromatin architecture may have significantly impacted the evolution of transcriptional regulation by increasing regulatory complexity (the combination of enhancer and promoter contacts)^53^. Taken together, these findings indicate that the evolution of gene regulation is strongly correlated with the 3D chromatin organization, especially intra-TAD interactions involving divergent CTCF sites. Interestingly, the divergent CTCF sites that primarily reside at TAD interiors have been spread through genomes by retrotransposition of B2 SINE transposable elements, which contain CTCF motifs in their consensus sequence^19,20^. Regarding these facts, our data illustrate that the chromatin remodeler, CHD4, conceals the B2 SINE-originated divergent/inappropriate CTCF-binding sites by regulating chromatin accessibility, thereby preventing three events: (1) aberrant CTCF recruitments (2) aberrant CTCF binding-associated changes in local TADs (separating and weakened TADs) and (3) overexpression of B2 SINE transcripts in mESCs.

The previous chromatin remodeler studies mainly focused on examining the ATP catalysis-based nucleosome sliding *in vitro*^54–57^. More recent reports have investigated the distribution of chromatin remodelers throughout the genome^58^, especially at promoters-TSS^26^, and their functional role in diverse cellular processes^23–26^. Such studies showed that CHD1 plays a critical role in maintaining the open chromatin structure by preventing heterochromatinization and is also required for pluripotency in mESCs^23^. In addition, various chromatin remodelers (CHD1, CHD2, CHD4, CHD6, CHD8, CHD9, BRG1, and EP400) have been shown to act together at specific nucleosomes near promoters and regulate transcription in mESCs^26^. CHD4 is also known to be part of the protein network underlying pluripotency (PluriNetwork)^59^ and critically ensures mESC identity by regulating pluripotency- and differentiation-associated genes^60^. However, the detailed/genome-wide molecular mechanisms through which chromatin remodelers function in mESCs have remained elusive, especially given that the prior studies have mainly focused on chromatin remodelers in transcribed regions, which account for only about 2% of the mammalian genome.

Regarding other aspects than transcribed regions, several reports showed that chromatin remodelers are closely associated with CTCF. Previously, RNAi-mediated knockdown of *CHD8* revealed that CHD8 interacts with CTCF and is required for the CTCF-dependent insulator function at specific CTCF-binding sites (*H19*, control region of *β-globin* and promoter regions of *BRCA1* and *c-myc* genes) in HeLa and Hep3B cells^61^. In addition, RNAi-mediated knockdown of *SNF2H* (*SMARCA5*) uncovered that SNF2H promotes CTCF binding (maintains CTCF occupancy) and organizes the nucleosome positioning adjacent to CTCF-binding sites but does not act as a CTCF’s loading factor in HeLa cells^62^. Interestingly, they also examined other chromatin remodelers, including CHD4, but did not detect major changes in CTCF occupancy and nucleosome occupancy/positions near CTCF-binding sites. These different results compared to ours may be caused by using different cell lines (mESC vs. HeLa) or measuring the changes in CTCF/Nucleosome occupancy/position as average line plots, which only exhibit the general/global changes rather than context-specific changes. Recently, conditional knockout of *Chd4* in granule neurons of mouse cerebellum revealed that CHD4 regulates chromatin accessibility and cohesin binding, thereby coordinates intra-domain loop strength and gene expression in mouse brain^63^. Furthermore, knockout of *Adnp*, which forms ChAHP complex with CHD4 and HP1β/γ^64^, in mESCs found that ChAHP complex, which mainly localized at less diverged B2 SINEs, prevents aberrant CTCF recruitments and maintaining TAD organizations^65^. Regarding these, the results obtained from *Adnp* knockout and our CHD4 depletion seem very similar. However, when comparing these two, the primary mechanism for maintaining proper CTCF binding is explicitly different. In the case of ADNP, the ChAHP complex competes with CTCF for common binding sequences (motifs), thereby preventing the aberrant CTCF bindings. In the ADNP study, the authors’ primary focus was ADNP; all of the data were obtained by the deletion of ADNP, not CHD4, and the functional role of CHD4 was not their major interest. On the other hand, our primary target is CHD4. Notably, we found that CHD4 directly regulates chromatin accessibility at putative/aberrant CTCF motifs (Group 1 CTCF-binding sites), concealing these sites as H3K9me3-enriched heterochromatic regions, thereby preventing aberrant CTCF bindings. When CHD4 is intact, the aberrant CTCF-binding sites are concealed by nucleosomes, exhibiting the closed chromatin states; these were confirmed by the enriched nucleosomes (Fig. 3c-e), core histones (Fig. 4, Supplementary Fig. 3H), constitutive heterochromatin marker H3K9me3 (Fig. 3f, Supplementary Fig. 3H), and depleted DNaseI and ATAC-seq signals (Fig. 3c-e). Upon CHD4 depletion, the concealed aberrant CTCF-binding sites become accessible; these were confirmed by the fact that enriched nucleosomes (Fig. 3c-e), core histones (Fig. 4), and H3K9me3 (Fig. 3f) are lost, while ATAC-seq signals are increased (Fig. 3c-e). Furthermore, we performed temporal depletion/restoration of CHD4 to determine the order of events (Fig. 4, Supplementary Fig. 4); CHD4 first assembles core histones at the aberrant CTCF-binding sites and thereby prevents aberrant CTCF recruitments. We also performed rescue experiments by transfecting ectopic CHD4 after CHD4 depletion and confirmed that this aberrant CTCF binding is a direct effect caused by CHD4 depletion (Fig. 5, Supplementary Fig. 5). Lastly, while the ChAHP study found that ADNP specifically binds to less diverged B2 SINEs, our data showed direct evidence that CHD4 regulates chromatin accessibility at B2 SINE-embedded aberrant CTCF-binding sites (Group 1 sites); CHD4 depletion triggered the overexpression of B2 SINE transcripts in mESCs (Fig. 6, Supplementary Fig. 6). Thus, there are significant differences in major mechanisms for regulating CTCF bindings involving ADNP (ChAHP) and CHD4.

Based on our findings, we propose a mechanism wherein the ATP-dependent chromatin remodeler, CHD4, maintains the lower-level chromatin structure (including chromatin accessibility and nucleosome positioning/occupancy) and prevents the aberrant recruitment of CTCF, thereby safeguarding the appropriate CTCF bindings and associated higher-order TAD organizations by suppressing aberrant interactions at TAD interiors (Fig. 7). Together, we uncovered how the primary structure of chromatin could facilitate higher-order 3D chromatin organization in mESCs. Collectively, our findings provide a novel perspective on the functional role of chromatin remodelers in mESCs, thereby adding an extra layer of information in stem cell biology and greatly facilitating our understanding of chromatin remodelers in embryonic stem cells.

## Acknowledgements

We thank Taemmok Kim and Prof. Keunsoo Kang for assistance with analyzing the NGS data; Kwang-Beom Hyun and Prof. Jaehoon Kim for purification of Chd4 protein using the Bac-to-Bac Baculovirus expression system (Full-length cDNA of mouse Chd4 was purchased from Open Biosystems, MMM1013-202770503). This research was supported by grants from the Stem Cell Research Program (2012M 3A9B 4027953) through the National Research Foundation of Korea (NRF) funded by the Ministry of Science and ICT. This work was also supported by a National Research Foundation of Korea grant funded by the Korean Government (2018R1A5A1024261, SRC). The authors declare that no competing interests exist.

## Author Contributions

S. Han, H. Lee and D. Lee designed the concepts and experiments. S. Han analyzed most of the NGS data. H. Lee performed most of the experiments. A. J. Lee performed the Hi-C experiments under the guidance of I. Jung. S.-K. Kim performed the GRO-seq experiments under the guidance of T.-K. Kim. S. Han and H. Lee wrote the manuscript under the mentorship and technical supervision of G. Y. Koh and D. Lee.

## Methods

### Mouse ES cell culture

E14Tg2a mESCs were maintained under feeder-free conditions. Briefly, cells were cultured on gelatin-coated cell culture dishes in an mESC culture medium consisting of Glasgow’s minimum essential medium (GMEM) containing 10% knockout serum replacement, 1% non-essential amino acids, 1% sodium pyruvate, 0.1 mM β-mercaptoethanol (all from Gibco), 1% FBS, 0.5% antibiotic-antimycotic (both from Hyclone) and 1,000 units/ml LIF (ESG1106, Millipore). mESCs were maintained at 37°C with 5% CO_2_ in humidified air.

### RNA interference

The siRNAs against *EGFP* and *Chd4* were synthesized and annealed by ST Pharm (Seoul, Korea). Their sequences are presented in Supplementary Table 5. mESCs were transfected with 50 nM of the indicated siRNA using DharmaFECT I (T-2001-03, Dharmacon) according to the manufacturer’s protocol. Briefly, mESCs were seeded to 6-well plates. One day later, 50 nM of siRNAs and DharmaFECT reagent were diluted in Opti-MEM (Gibco), incubated separately at 25°C for 5 min, and then mixed together. The mixtures were incubated at 25°C for 20 min and added to the mESC cultures. The culture medium was replaced after 24 hours. Transfected mESCs were harvested at 48 hours after transfection, and knockdown efficiency was analyzed by real-time quantitative polymerase chain reaction (RT-qPCR).

### RNA purification and reverse transcription

Total RNAs were purified from mESCs using the TRIzol reagent (Invitrogen) according to the manufacturer’s protocol. Briefly, mESCs cultured in 6-well plates were harvested and homogenized with 1 ml of TRIzol reagent. Chloroform (200 μl/sample) was added, and the samples were mixed vigorously by hand for 15 sec and incubated at 25°C for 2 min. The mixtures were centrifuged at 12,000 rpm for 15 min at 4°C, and 500 μl of each aqueous phase was transferred to a new Eppendorf tube and mixed with the same volume of isopropanol. The mixtures were incubated at 25°C for 10 min to precipitate total RNAs. The samples were centrifuged at 12,000 rpm for 10 min at 4°C, washed with 75% ethanol, and centrifuged again at 10,000 rpm for 5 min at 4°C. The RNA pellets were dried and dissolved in RNase-free water, and 1 μg of DNase-treated total RNA was applied for cDNA synthesis using an Improm-II Reverse transcription system (A3802, Promega) according to the manufacturer’s protocol. For analysis of retrotransposon expression, the cDNA synthesis step was primed with random hexamers.

### Real-time quantitative polymerase chain reaction (RT-qPCR)

The generated cDNAs were amplified using a BioFact Real-time PCR kit (BIOFACT, Daejeon, Korea) according to the manufacturer’s manual. The primer sequences used for RT-qPCR are presented in Supplementary Table 6. Briefly, 20-μl reactions containing 1x EvaGreen, 10 mM tetraethylammonium chloride, and 10 pmol of primers were analyzed with a CFX96 system (Bio-Rad) under the following conditions: 95°C for 12 min (initial melting), followed by 40 cycles of 95°C for 20 sec (denaturation), 57°C for 30 sec (annealing), and 72°C for 30 sec (extension). The relative expression levels of Chd family members and retrotransposons were quantified with respect to those of β-actin and the 28S rRNA, respectively.

### Generating CHD4-mAID mESCs

We employed the auxin-inducible degron (AID) system to *Chd4* gene and generated the stable cell line for CHD4-mAID E14Tg2a mESCs as previously described^16,30^. Briefly, we first generated OsTIR1 parental cell line (Supplementary Fig. 1G, H) by transfecting pEN396-pCAGGS-Tir1-V5-2A-PuroR (*Tigre* donor, #92142, Addgene) and pX330-EN1201 (spCas9nuclease with *Tigre* sgRNA, #92144, Addgene) to E14Tg2a mESCs. After confirming the successful generation of OsTIR1 parental cell (Supplementary Fig. 1H), we cloned *Chd4* donor based on pMK293-mAID-mCherry2-Hygro (#72831, Addgene) and also cloned spCas9nuclease with *Chd4* sgRNA based on PX458-pSpCas9(BB)-2A-GFP (#48138, Addgene). Then, we generated CHD4-mAID E14Tg2a by transfecting these two vectors to OsTIR1 parental cell. Degradation of AID-tagged CHD4 was induced by the addition of 500 μM 3-Indoleacetic acid (I2886, Sigma).

### CUT&RUN

CUT&RUN assays were performed as previously described^29,66^, with minor modification. Briefly, 4 million mESCs were harvested and washed with 1.5 ml Wash buffer (20 mM HEPES, pH 7.5, 150 mM NaCl, 0.5 mM Spermidine) three times. Cells were bound to activated Concanavalin A-coated magnetic beads (at 25°C for 10 min on a nutator), then permeabilized with Antibody buffer (Wash buffer containing 0.05% Digitonin and 4 mM EDTA). The bead-cell slurry was incubated with 3 μl relevant antibody (see below) in a 150 μl volume at 25°C for 2 hours on a nutator. After two washes in 1 ml Dig-wash buffer (Wash buffer containing 0.05% Digitonin), beads were resuspended in 150 μl pAG/MNase and incubated at 4°C for 1 hour on a nutator. After two washes in 1 ml Dig-wash buffer, beads were gently vortexed with 100 μl Dig-wash buffer. Tubes were chilled to 0°C for 5min and ice-cold 2.2 mM CaCl_2_ was added while gently vortexing. Tubes were immediately placed on ice and incubated at 4°C for 1 hour on a nutator, followed by addition of 100 μl 2xSTOP buffer (340 mM NaCl, 20 mM EDTA, 4 mM EGTA, 0.05% Digitonin, 0.1 mg/ml RNase A, 50 μg/ml glycogen) and incubated at 37°C for 30min on a nutator. Beads were placed on a magnet stand and the liquid was removed to a fresh tube, followed by addition of 2 μl 10% SDS and 2.5 μl proteinase K (20mg/ml) and incubated at 50°C for 1 hour. DNA was extracted using phenol chloroform as described at https://www.protocols.io/view/cut-amp-run-targeted-in-situ-genome-wide-profiling-zcpf2vn. CUT&RUN libraries were prepared using a Accel-NGS 2S Plus DNA Library Kit (21024, SWIFT), according to the manufacturer’s guidelines. The libraries were then sequenced using an Illumina Novaseq 6000 platform. The libraries were generated from two sets of biological replicates.

### *In-situ* Hi-C

*In-situ* Hi-C was performed as previously described^6^. Briefly: 1.5 million mESCs were crosslinked with 1% formaldehyde; nuclei were isolated; chromatin was digested with MboI (R0147, NEB); 5’ ends were filled by incorporation of biotinylated-dCTP (19524-016, Life Technologies); proximity ligation, reverse-crosslinking, and DNA shearing were performed; biotinylated junctions were isolated with streptavidin beads (65601, Life Technologies); and DNA libraries were prepared. Each Hi-C library was amplified for six cycles and then subjected to deep sequencing on a Hiseq4000 Illumina platform. The libraries were generated from two sets of biological replicates.

### ChIP-seq

ChIP assays were performed as previously described^67^, with minor modification. Briefly, mESCs were washed twice with PBS and incubated with 1% formaldehyde for 15 min at 25°C. This crosslinking was quenched with 125 mM glycine at 25°C for 5 min, and the cells were harvested with cold PBS and suspended in SDS lysis buffer (50 mM Tris-HCl, pH 8.0, 1% SDS, 10 mM EDTA). The chromatin was sheared to mono- or dinucleosome sizes using a focused ultrasonicator (Covaris S220). The sonicates were incubated overnight with the relevant antibodies (see below) and Protein A, G sepharose (17-1279-03 and 17-0618-05, GE Healthcare) at 4°C on a nutator. The immune complexes were washed for 10 min each with the following wash buffers: low-salt wash buffer (20 mM Tris-HCl, pH 8.0, 150 mM NaCl, 0.1% SDS, 1% Triton X-100, 2 mM EDTA), high-salt wash buffer (20 mM Tris-HCl, pH 8.0, 500 mM NaCl, 0.1% SDS, 1% Triton X-100, 2 mM EDTA), and LiCl wash buffer (10 mM Tris, pH 8.0, 250 mM LiCl, 1% NP-40, 1% sodium deoxycholate, 1 mM EDTA). The immune complexes were further washed twice with TE buffer (10 mM Tris-HCl, pH 8.0, 1 mM EDTA). Finally, the immune complexes were eluted with elution buffer (1% SDS, 0.1 M NaHCO_3_) and reverse-crosslinked overnight at 68°C. The immunoprecipitated DNA was treated with proteinase K and RNase A and recovered by phenol-chloroform-isoamyl alcohol precipitation. ChIP-seq libraries were prepared using a NEXTflex ChIP-seq kit (5143-02, Bioo Scientific) according to the manufacturer’s guidelines. The libraries were then sequenced using an Illumina HiSeq 2500 platform. The libraries were generated from two sets of biological replicates.

### Antibodies

Antibodies against Chd4 (ab70469, Abcam), CTCF (ab37477, Abcam), and β-actin (SC-47778, Santa Cruz Biotech) were used for immunoblotting. Antibodies against Chd4 (ab70469, Abcam and in-house generated) and IgG (12-371, Millipore) were used for immunoprecipitation. Antibodies against CTCF (07-729, Millipore), H3 (ab1791, Abcam), H3K9me3 (ab8898, Abcam), H3K4me1 (in-house generated), H3K4me3 (in-house generated), H3K27ac (ab4729, Abcam), and H3K27me3 (07-449, Millipore) were used for ChIP-seq.

### Immunoblotting

E14Tg2a mESCs were transfected with siRNAs against *EGFP* and *Chd4*. The transfected cells were harvested, washed with cold PBS, and lysed with EBC buffer (50 mM Tris-HCl, pH 8.0, 300 mM NaCl, 0.5% NP-40, 1 mM PMSF). The lysates were boiled for 5 min with SDS sample buffer, resolved by SDS-PAGE, and subjected to immunoblotting.

### ATAC-seq

ATAC-seq libraries were prepared as previously described^68,69^, with minor modification. Briefly, 50,000 mESCs were harvested, washed with cold PBS, lysed with cold lysis buffer, and immediately centrifuged. The nuclear pellets were resuspended in 25 μl of 2X tagmentation reaction buffer (10 mM Tris, pH 8.0, 5 mM MgCl_2_, 10% dimethylformamide), 23 μl of nuclease-free water, and 2 μl of Tn5 transposase (in-house generated), and incubated at 37°C for 30 min. The samples were then immediately purified using a QIAquick PCR purification kit (28106, Qiagen). The libraries were pre-enriched for five cycles using the KAPA HiFi Hotstart ready mix (KK2601, Kapa Biosystems), and the threshold cycle (Ct) was monitored using qPCR to determine the additional enrichment cycles, which were then applied. The final libraries were purified again with a QIAquick PCR purification kit, and sequenced using an Illumina HiSeq 2500 platform. The libraries were generated from two sets of biological replicates.

### Micrococcal nuclease digestion and MNase-seq

Cells cultured in 60Ф culture dishes were washed with PBS and incubated with modified PBS (150 mM NaCl, 0.02% Tween-20, 0.02% Triton X-100) at 25°C for 5 min. The cells were then fixed with 1% formaldehyde at 25°C for 10 min, quenched with 125 mM glycine at 25°C for 5 min, washed, and suspended with lysis buffer (18% Ficoll-400, 10 mM KH_2_PO_4_, 10 mM K_2_HPO_4_, 1 mM MgCl_2_, 250 nM EGTA). The lysates were centrifuged at 12,000 rpm for 40 min at 4°C, and the pelleted chromatin was resuspended in A buffer (10 mM Tris-Cl, pH 7.4, 150 mM NaCl, 5 mM KCl, 1 mM EDTA). CaCl_2_ (5 mM) was added to the samples, which were immediately digested with MNase (M0247S, NEB) at 37°C for 60 min. The reaction was inactivated with 1% SDS and 500 mM EDTA, and reverse-crosslinking was performed overnight at 68°C. Proteinase K and RNase were added sequentially to the samples, and DNAs were recovered using phenol-chloroform-isoamyl alcohol extraction. The MNase-digested DNAs were separated on a 1.5% agarose gel, and mono-nucleosomal DNAs were extracted using a QIAEX II gel extraction kit (20021, Qiagen) according to the manufacturer’s manual. The purified mono-nucleosomal DNAs were subjected to sequencing using a TruSeq DNA library prep kit (FC-121-2001, Illumina). The final libraries were sequenced using an Illumina HiSeq 2500 platform. The libraries were generated from two sets of biological replicates.

### Total RNA-seq and mRNA-seq

DNase-treated total RNAs were purified from mESCs and used for total RNA-seq. Total RNA-seq libraries were prepared using a TruSeq stranded RNA kit (RS-122-2301, Illumina) according to the manufacturer’s manual. For mRNA-seq library preparation, mRNAs were isolated from total RNA using a Magnetic mRNA isolation kit (S1550S, NEB), and libraries were prepared using a NEXTflex Rapid directional RNA-seq kit (5138-08, Bioo Scientific). The libraries were sequenced using an Illumina HiSeq 2500 platform. The libraries were generated from two sets of biological replicates.

### Immunoprecipitation

E14Tg2a mESCs were harvested, washed with cold PBS, and lysed with EBC buffer (50 mM Tris-HCl, pH 8.0, 150 mM NaCl, 0.5% NP-40, 1 mM PMSF). The lysates were incubated for 1 hour with the indicated antibodies and Protein A, G sepharose (17-1279-03 and 17-0618-05, GE Healthcare) at 4°C with agitation. The immune complexes were washed three times with EBC buffer (50 mM Tris-HCl, pH 8.0, 150 mM NaCl, 0.5% NP-40, 1 mM PMSF). The lysates were boiled for 5 min with SDS sample buffer, resolved by SDS-PAGE, and subjected to immunoblotting.

### RNA immunoprecipitation (RIP) assays

RIP assays were conducted as previously described^70,71^, with minor modification. Briefly, cultured mESCs were crosslinked with 1% formaldehyde for 15 min at 25°C and quenched with 125 mM glycine. The cells were harvested, washed with cold PBS, and then lysed with RIPA buffer (50 mM Tris-Cl, pH 8.0, 150 mM NaCl, 0.05% SDS, 1 mM EDTA, 1% NP-40, and 0.05% sodium deoxycholate) containing an RNase inhibitor (M007L, Enzynomics). The samples were sonicated, treated with DNase I (18068-015, Invitrogen) for 10 min at 37°C, and cleared by centrifugation. The extracts were bound with anti-Chd4 (ab70469, Abcam) for 2 hours at 4°C in the presence of RNase inhibitor. Immune complexes were further incubated with Dynabeads Protein A and G (10001D, 10004D, Invitrogen) for 1 hour at 4°C. The immune complexes were then washed for 10 min each with the following wash buffers: low-salt wash buffer (20 mM Tris-HCl, pH 8.0, 150 mM NaCl, 0.1% SDS, 1% Triton X-100, 2 mM EDTA), high-salt wash buffer (20 mM Tris-HCl, pH 8.0, 500 mM NaCl, 0.1% SDS, 1% Triton X-100, 2 mM EDTA), and LiCl wash buffer (10 mM Tris, pH 8.0, 250 mM LiCl, 1% NP-40, 1% sodium deoxycholate, 1 mM EDTA). The immune complexes were further washed twice with TE buffer (10 mM Tris-HCl, pH 8.0, 1 mM EDTA), eluted, and reverse-crosslinked with 400 mM NaCl for 2 hours at 65°C. The RNAs were purified from the eluates using an RNA Clean & Concentrator™-5 kit (R1015, Zymo Research) according to the manufacturer’s manual, and then incubated again with DNase I (18068-015, Invitrogen) for 30 min at 37°C. Finally, DNase I was inactivated and removed using DNA-free^TM^ kit (AM1906, Invitrogen) according to the manufacturer’s manual.

### *In vitro* transcription and biotinylation of RNAs

The plasmids used for *in vitro* transcription were constructed as previously described^51^, with minor modification. The primer sequences used for *in vitro* transcription are presented in Supplementary Table 7. Briefly, we used the pUC vector to construct pUC-T7-yGAL1, pUC-T7-EGFP1/2, and pUC-T7-Nanog. The indicated RNAs were transcribed *in vitro* using a TranscriptAid T7 High Yield Transcription kit (K0441, Thermo Scientific) and then purified using an RNA Clean & Concentrator™-5 kit according to the manufacturer’s manual. The purified RNAs were resolved on 5% denaturing polyacrylamide mini gels at 70V, 4°C for 2 hours. The gels were cut for size selection and appropriately sized fragments were eluted with elution buffer (10 mM Tris-HCl, pH 7.5, 300 mM NaCl, 0.1% SDS, 1 mM EDTA) overnight at 4°C with agitation. The eluted RNAs were purified using an RNA Clean & Concentrator™-5 kit. The purified RNAs of this step were used for the nucleosome-sliding assay. For other experiments, the purified RNAs were labeled with biotin using a Pierce RNA 3’ End Biotinylation kit (20160, Thermo Scientific) according to the manufacturer’s manual. The biotinylated RNAs were purified using an RNA Clean & Concentrator™-5 kit and then resolved 5% on a denaturing polyacrylamide mini gel at 70V, 4°C for 2 hours. The gels were cut for size selection and eluted with elution buffer (10 mM Tris-HCl, pH 7.5, 300 mM NaCl, 0.1% SDS, 1 mM EDTA) overnight at 4°C with agitation. The eluted biotinylated RNAs were purified once more using an RNA Clean & Concentrator™-5 kit, and then subjected to EMSA.

### Electrophoretic mobility shift assay (EMSA)

EMSA was carried out as previously described^71^, with minor modifications. Briefly, the indicated amount of purified Chd4 protein was incubated with 5 nM of biotinylated RNAs in binding buffer (50 mM Tris-Cl, pH 7.5, 100 mM NaCl, 10 mM β-mercaptoethanol, and 5% glycerol) for 40 min at 25°C. Bound protein-RNA complexes were resolved on 5% native polyacrylamide mini gels at 70V, 4°C for 2 hours. The complexes were transferred to an Amersham Hybond-N+ nylon membrane (RPN303B, GE Healthcare), UV-crosslinked, and visualized using a Chemiluminescent Nucleic Acid Detection Module kit (89880, Thermo Scientific) according to the manufacturer’s manual.

### Nucleosome reconstitution and nucleosome-sliding assay

Nucleosome reconstitution was performed as previously described^72^. Briefly, two types (lateral and central forms) of a 147-bp DNA fragment were amplified from pGEM-3z/601, and *X. laevis* core histones were purified using bacterial expression systems and nickel-affinity chromatography. The core histones and DNA fragments were mixed at a 1:1 ratio in initial buffer (10 mM HEPES, pH 7.9, 1 mM EDTA, 5 mM DTT, 0.5 mM PMSF), brought to 2 M NaCl with 2 μg BSA in each 10 μl reaction, and incubated for 15 min at 37°C. The reaction was serially diluted with 3.6 μl, 6.7 μl, 5 μl, 3.6 μl, 4.7 μl, 6.7 μl, 10 μl, 30 μl, and 20 μl of initial buffer, and each dilution was incubated for 15 min at 30°C. Lastly, the reaction was diluted with 100 μl of final buffer (10 mM Tris-HCl, pH 7.5, 1 mM EDTA, 0.1% NP-40, 5 mM DTT, 0.5 mM PMSF, 20% glycerol, 100 μg/ml BSA) and incubated for 15 min at 30°C. The lateral and central nucleosome forms were reconstituted and used for the nucleosome-sliding assay.

For the nucleosome-sliding assay, reconstituted nucleosomes (75 ng, 0.36 pmol) were incubated with purified Chd4 (50 ng, 0.23 pmol), varying concentrations of purified RNAs (0 pmol, 0.1 pmol, 0.2 pmol, 0.5 pmol, and 1 pmol), and ATP (0.1 pmol) in sliding buffer (50 mM KCL, 20 mM HEPES, pH 7.9, 2 mM DTT, 0.5 mM PMSF, 0.05% NP40, 10% glycerol, 100 μg/ml BSA, 10 mM MgCl_2_) for 45 min at 30°C. The reaction was stopped by the addition of 500 ng of plasmid (size > 10 kb) and 0.1 pmol of ATP-γ-S tetralithium salt (10102342001-Roche, Sigma) followed by incubation for 30 min at 30°C. The samples were resolved on 5% native polyacrylamide mini gels at 65V, 4°C for 2.5 hours.

### Global run-on (GRO)-seq

Circularized GRO-seq was performed as previously described^73,74^. Briefly, around 10 million nuclei per sample were purified and used for global run-on and base hydrolysis, which were carried out as previously described^75^. BrU-labeled nascent RNAs were immunoprecipitated twice with anti-BrdU antibody-conjugated agarose (SC-32323ac, Santa Cruz Biotech). Between the two immunoprecipitations, the BrU-precipitated RNAs were subjected to polyA tailing with poly(A)-polymerase (P7460L, Enzymatics). The RNAs were then subjected to first-strand cDNA synthesis using Superscript III Reverse Transcriptase (18080-044, Invitrogen) and the oNTI223 RT primer (IDT). Excess RT primers were removed with exonuclease I (M0293S, NEB), and the cDNAs were size-selected (120-220 nt) in a 6% polyacrylamide TBE-urea gel. The selected cDNAs were subsequently circularized using CircLigase (CL4111K, Epicentre) and relinearized with ApeI (M0282S, NEB). The single-stranded DNA templates were amplified using Phusion High-Fidelity DNA Polymerase (M0530S, NEB) and Illumina TruSeq small-RNA sample barcoded primers. The generated PCR products were isolated and size-selected (190-290 bp) by electrophoresis on a 6% native polyacrylamide TBE gel. The final libraries were sequenced on an Illumina HiSeq 2500 platform. The libraries were generated from two sets of biological replicates.

### Gene Ontology analysis

To find the gene ontology terms of differentially expressed genes (DEGs), we used ConsensusPathDB^76^.

### Data processing and analysis

The results of our ChIP-seq, MNase-seq, ATAC-seq, and GRO-seq analyses were aligned to the mouse genome (UCSC mm10) using Bowtie2 (version 2.2.9) with default parameters^77^. The results of our mRNA-seq and total RNA-seq analyses were aligned to the mouse genome (UCSC mm10) using STAR (version 2.5.2a) with default parameters^78^. For the ChIP-seq, ATAC-seq, MNase-seq, mRNA-seq, total RNA-seq, and GRO-seq analyses, we used MACS2^79^ to convert the aligned BAM files into bedGraph files, and used MACS2 (--SPMR) to normalize the data with respect to the total read counts. ChIP-seq data were further normalized with respect to the input. Then, we used bedGraphToBigWig^80^ to convert the bedGraph files into bigWig files. The bigWig files were used as input files for bwtool (matrix option) to quantify the intensity (e.g., using heatmaps) of the relevant sequencing data^81^. Further analyses were performed using in-house scripts, MACS2^79^, and HOMER^82^. All of our raw data (fastq files) were confirmed to be of good quality using FastQC (http://www.bioinformatics.babraham.ac.uk/projects/fastqc/).

For our ChIP-seq analysis, we used MACS2 (callpeak option, *P*-value < 0.005). We then double checked our findings using HOMER (findPeaks option) to investigate specific regions that were enriched with the relevant histone modifications and factors^79,82^. Both MACS2 and HOMER compare ChIP data with Input files and return significantly enriched sites. The ChIP-seq data were also subjected to motif analysis, which was performed using the HOMER findMotifsGenome.pl option, and analyses of the genomic contents within CTCF-binding sites were performed using the HOMER annotatPeaks.pl option^82^. The bigWig files (Chd4KD/Control or IP/Input) were divided using deepTools (bigwigCompare option, --ratio=log2)^83^.

For our mRNA-seq and total RNA-seq analyses, we used Cufflinks (Cuffdiff option, fr-firststrand). We then double checked our findings with HOMER (analyzeRNA.pl and analyzeRepeats.pl option)-based quantification of expression levels^82,84,85^. For our analysis of SINEs, we built a STAR-index exclusively containing the previously identified 1.5 million SINE copies (RepeatMasker in UCSC table browser), and performed STAR and Cufflinks analyses as described above.

For our GRO-seq analysis, we used HOMER (findPeaks option, -style groseq) to examine *de novo*-produced SINE transcripts (Extended Data Fig. 8e)^82^.

Box plots, volcano plots, and other plots were drawn with R (ggplot2)^86^ and heatmaps were drawn with Java TreeView^87^. The examples of our genome-wide data were visualized using the Integrative Genomics Viewer (igv)^88^.

For our Hi-C analysis, each raw sequencing data set was aligned to mm10 using BWA-mem with default parameters. Chimeric reads that spanned ligation sites were processed using an in-house script; such reads are the result of the ligation chemistry used during Hi-C library construction, and are not properly processed by the paired-end BWA-mem command. The read-pairs were merged together as paired-end-aligned bam files, and PCR duplicates were removed with Picard (http://broadinstitute.github.io/picard). To remove multiple sources of intrinsic biases relevant to the raw Hi-C data, we applied negative binomial model-based implicit normalization approaches and created a 20-kb resolution normalized interaction matrix. To enable a robust comparative analysis and compensate for differences in library size, we performed quantile-normalization between Control (E14Tg2a mESCs transfected with *siEGFP*) and Chd4KD (E14Tg2a mESCs transfected with *siChd4*) samples for each chromosome. Lastly, we used PCA (principal component analysis) to obtain PC1 (first principal components, equivalent to the first eigenvectors) values for Control and Chd4KD cells (HOMER, runHiCpca.pl).

### Public data acquisition

The publically released ChIP-seq data were downloaded from NCBI GEO DataSets and the ENCODE Consortium. These data were downloaded as sra or fastq formats, and sra files were converted to fastq files using the SRA Toolkit (https://trace.ncbi.nlm.nih.gov/Traces/sra/sra.cgi?view=software). Thus, we analyzed the public datasets and our own results using the same methods. Details of the analyzed data are summarized in Supplementary Table 1.

---

**Supplementary Fig. 1**

**CHD4 and CTCF are closely linked, and CHD4 depletion marginally affects the global 3D chromatin organizations.**

**A** RT-qPCR analysis of *Chd* family members in various mouse cell lines, including mouse embryonic stem cells (mESCs), mouse embryonic fibroblasts (MEFs), and NIH3T3 cells. Error bars denote the standard deviation obtained from three biological replicates.

**B** Motif analysis of CHD4-binding sites and random binding sites, as carried out using MEME. CHD4^1^ (in-house generated antibody) and CHD4^2^ (Abcam antibody) denote CUT&RUN data using different antibodies against CHD4. Random^1^ and random^2^ have the same numbers and the median length of binding sites as CHD4^1^ and CHD4^2^, respectively, but the peak positions were randomly selected. See also Supplementary Table 2.

**C, D** Heatmaps of CHD4 aligned at 2,153 TADs^4^ (**C**) and 23,726 insulated neighborhoods^8^ (**D**). All heatmaps were sorted in ascending order by the length of the TAD or insulated neighborhood. CHD4^3^ (de Dieuleveult, M., et al. Nature. 2016.), CHD4^4^ (Luo, Z., et al. Mol Cell. 2015.), and CHD4^5^ (Whyte, W.A., et al. Nature. 2012.) denote ChIP-seq data obtained from other studies. See also Supplementary Table 1.

**E** RT-qPCR analysis of *Chd4* following siRNA-mediated knockdown of *Chd4* in mESCs. Gray and red bars indicate cells transfected with *siEGFP* (Control) and *siChd4* (Chd4KD), respectively. Error bars denote the standard deviation obtained from three biological replicates.

**F** Immunoblots of CHD4. α-Tubulin was detected as a loading control.

**G** To create E14Tg2a-CHD4-mAID cells (CHD4-mAID), we first integrated the V5-tagged OsTIR1 gene at the *Tigres* locus (OsTIR1 parental cell), and then both *Chd4* alleles were tagged with auxin-inducible degrons and mCherry2 reporter (CHD4-mAID cell).

**H** Immunoblots of OsTIR1-V5.

**I** RT-qPCR analysis of target regions within mAID (two red arrows) provided five candidates (red) of CHD4-mAID cells.

**J** Immunoblots of CHD4 and mCherry2 after auxin treatment in E14Tg2a wild-type (WT) and CHD4-mAID cells. α-Tubulin was detected as a loading control. See also Supplementary Fig. 4B.

**K** Genomic DNA PCR analysis of two types of target regions (two red arrows) provided the homozygous (red) and heterozygous carrying CHD4-mAID.

**L** RT-qPCR analysis of target regions (two red arrows) confirmed the homozygous CHD4-mAID cells.

**A, E, I, L** The expression levels were normalized with respect to that of β-actin.

**M** Scatter plots showing high reproducibility among the replicates of all Hi-C samples, as assessed using contact probability based on Pearson correlation.

**N** Relative contact probability (RCP) plots showing contact frequency relative to genomic distance.

**O** Scatter plots comparing genome-wide PC1 (first principal components, equivalent to the first eigenvectors) values between Chd4KD with Control cells (left) and +Aux with –Aux cells (right). The correlation coefficient (R^2^) was calculated. Gray dashed lines represent linear regression lines.

**P** Stacked bar graphs showing the percentage of each compartment type. Red and green boxes indicate static A and B compartments, respectively, while blue boxes represent changes in compartment domains (A to B or B to A) upon CHD4 depletion.

**Supplementary Fig. 2**

**Chd4 plays an important role in maintaining the 3D chromosome architectures by preventing aberrant CTCF bindings.**

**A** Venn diagrams representing the overlap of weakened and separating TADs or strengthened and merging TADs upon CHD4 depletion.

**B, C** Example of weakened TADs upon *Chd4* knockdown (left) and separating TADs upon CHD4 depletion (right) (**B**). Example of weakened TADs upon CHD4 depletion (**C**). Hi-C interactions (heatmaps), TADs (green lines and black triangles), insulation heatmaps, CTCF ChIP-seq peaks (red and blue triangles indicate + and – directionalities of CTCF motifs), and genes are shown in Control, Chd4KD, -Aux, and +Aux cells. Black arrows indicate the region where 3D chromatin architectures are severely disrupted upon CHD4 depletion. The zoom-in CTCF ChIP-seq data represent the gained CTCF ([**B**], 1*; [**C**], 2* and 4*) which resides at a newly-established border of separating TADs or regions displaying potential border with decreased insulations within weakened TADs (dotted boxes) upon CHD4 depletion.

**D** Box plots showing the TAD sizes in mega-base (mb) scale. The horizontal line and the white rhombus in the box denote the median and mean, respectively. *P*-values were calculated using the Wilcoxon rank sum test.

**E, F** Aggregate peak analysis (APA) showing the Hi-C contacts (top) and differential interactions (bottom) of CHD4 depleted cells compared to Control (-Aux) cells at Control- or –Aux-specific peaks (**E**) and Chd4KD- or +Aux-specific peaks (chromatin loops) (**F**).

**G, H** Bar graphs showing the number of peaks (**G**) and specific peaks (**H**).

**I** Aggregate peak analysis (APA) showing the Hi-C contacts (top) and differential interactions (bottom) of CHD4 depleted cells compared to Control (-Aux) cells at gained CTCF-associated peaks (chromatin loops).

**E, F, I** Box plots (right) displaying the quantified contacts at these peaks. *P*-values were calculated using the Wilcoxon signed rank test.

**Supplementary Fig. 3**

**CHD4 does not affect the expression level of *CTCF* but instead controls the proper recruitment of CTCF by regulating chromatin accessibility at heterochromatic regions.**

**A, B** Heatmaps representing CHD4^2^, IgG^2^ CUT&RUN, CTCF ChIP-seq of Control and Chd4KD cells (**A**), and CTCF ChIP-seq of –Aux and +Aux cells (**B**) at CHD4 peaks (left), CHD4 peaks that coincide (top right), and not coincide with CTCF peaks (bottom right).

**C** Bar graphs showing the mRNA expression levels of *CTCF*.

**D** Immunoblots of CTCF. β-Actin was detected as a loading control.

**E, H-J** Heatmaps displaying CUT&RUN (CHD4^1,2^ and IgG^1,2^) and ChIP-seq signals as indicated on top. All heatmaps were aligned at 39,800 CTCF peaks (rows) and sorted in descending order by the relative CTCF intensity (Chd4KD/Control). 4,736 Group 1 (11.9%) represents CTCF-binding sites at which more CTCF were recruited in Chd4KD cells compared to Control cells (1.75-fold), while 35,064 Group 2 (88.1%) represents the remainder of the CTCF-binding sites, at which CTCF binding was unchanged or reduced in Chd4KD cells.

**F** Heatmaps displaying CTCF, RAD21, DNase I hypersensitive sites (DNase I), ATAC-seq signals (ATAC), the ratio of nucleosome-repelling sequences (AA-AAAA) or DNA bases (AT) to nucleosome-preferring sequences (GC-CG) or DNA bases (GC), and CTCF motifs as indicated on top. All heatmaps were aligned at 91,560 CTCF peaks (rows) and sorted in descending order by the relative CTCF intensity (+Aux/-Aux). 13,249 Group 1* (14.5%) represents CTCF-binding sites at which more CTCF were recruited in +Aux cells compared to -Aux cells (2-fold), while 78,311 Group 2* (85.5%) represents the remainder of the CTCF-binding sites, at which CTCF binding was unchanged or reduced in +Aux cells.

**G** Line plots showing average enrichments of CTCF and ATAC at Group 1* and Group 2* CTCF-binding sites. *P*-values were derived using Wilcoxon signed rank test (\**P* < 1×10^−100^; ns, not significant).

**E, F, H-J** Heatmaps that are not labeled as Control or Chd4KD represent wild-type mESCs. See also Supplementary Table 1.

**Supplementary Fig. 4**

**Temporal degradation/restoration reveals that CHD4 assembles core histones to prevent the aberrant CTCF binding at the Group 1 sites.**

**A** Examples of Group 1 sites, including representative results for CTCF and H3 across a time course of siRNA treatment.

**B** Immunoblots of CHD4 after various auxin treatment times in CHD4-mAID cells and E14Tg2a wild-type (E14.). α-Tubulin was detected as a loading control.

**C** Heatmaps displaying CTCF and H3 ChIP-seq of –Aux and +Aux followed by various auxin withdrawal times at all CTCF peaks in Control and Chd4KD cells as indicated on top. All heatmaps were aligned and sorted in the same way as described in Fig. 3c. See also Fig. 3c.

**D** Examples of Group 1* sites (defined as in Fig. 4k), including representative results for CTCF and H3 across a time course of auxin withdrawal.

**Supplementary Fig. 5**

**CHD4 binds with RNA, and this binding inhibits the nucleosome-sliding activity of CHD4 *in vitro*.**

**A** Immunoprecipitation of endogenous CHD4 from mESCs using the indicated antibodies. Whole-cell extracts were immunoprecipitated with two different CHD4 antibodies or IgG at 4°C. Immunoprecipitated samples were resolved by SDS-PAGE and immunoblotted using anti-CHD4^2^.

**B** The recombinant protein (FLAG-CHD4) used in this study was purified from a baculovirus expression system. The protein was resolved by SDS-PAGE and subjected to Coomassie blue staining.

**C** Independent biological replicates 2 (top) and 3 (bottom) of EMSAs for the *yGAL1* (left), *EGFP1* (middle), and *Nanog* (right) RNAs were performed with purified FLAG-CHD4. ‘Unbound RNA’ indicates free RNA, and ‘CHD4-RNA Complex’ indicates the binding of the purified protein to each RNA species.

**D** Nucleosome-sliding assays were performed using purified FLAG-CHD4 with the *yGAL1* (left), *EGFP1* (middle), and *EGFP2* (right) RNAs. ‘L’ and ‘C’ denote the lateral and central forms of nucleosomes, respectively. Nucleosome-sliding assays were performed with various RNA concentrations; 0.0, 0.1, 0.2, 0.5, and 1.0 nM of RNA was added in lanes 3, 4, 5, 6, and 7, respectively.

**E** Schematic representation of two methods for ectopic transfection of various FLAG-CHD4 constructs. Red arrows indicate the auxin treatment.

**F** Immunoblots of FLAG after two methods of ectopic transfection as described in Supplementary Fig. 5E.

**G** Immunoblots of FLAG after ++Aux method (Supplementary Fig. 5E, right) of ectopic transfection.

**F, G** α-Tubulin was detected as a loading control.

**H** Examples of Group 1* sites, including representative results for CTCF.

**Supplementary Fig. 6**

**CHD4 is required to suppress B2 SINE and may be involved in the proper recruitment of RNA Polymerase II (RNAPII) at the transcription start sites (TSSs) of pluripotent genes.**

**A** Examples of de-repressed B2 SINEs (GRO-seq, total RNA-seq, and mRNA-seq), aberrant CTCF bindings (CTCF ChIP-seq), increased chromatin accessibility (MNase-seq and ATAC-seq), and CHD4 enrichment at the Group 1 CTCF-binding sites in Chd4KD cells.

**B** Bar graphs showing the numbers of *de novo* SINE transcripts, as measured using GRO-seq.

**C** Volcano plots showing the differential expression of B2 SINE, as measured using total RNA-seq. B2 SINEs localized at the Group 1 CTCF-binding sites were analyzed with *P*-value (left) or without *P*-value (right). Blue and red dots indicate the expressions of SINEs that are significantly decreased and increased, respectively, upon *Chd4* knockdown (*P*-value ≤ 0.05 and log_2_ (Chd4KD/Control) ≤ −1 or ≥ +1).

**D** RT-qPCR analysis of retrotransposons in mESCs treated with α-amanitin for 6 hours before harvest. The expression levels were normalized with respect to that of the 28S rRNA, which is transcribed by RNA Pol I. Error bars denote the standard deviation obtained from three biological replicates.

**E** Result of GREAT (Genomic Regions Enrichment of Annotations Tool) analysis, which was used to identify a class of genes that are located near Group 1 CTCF-binding sites.

**F** Examples of RNAPII (ChIP-exo) at *Cox17* and *Snrpb* genes. Blue boxes highlight RNAPII at TSS.

**G** Bar graphs showing mRNA expression levels of RNAPII subunits.

**H** MA plots of exemplary genes in Fig. 6g and Supplementary Fig. 6F based on total RNA-seq (left) and mRNA-seq (right) data. Red dots indicate genes with abundant de-repressed B2 SINE near their TSSs, whereas gray dots indicate genes without de-repressed B2 SINE near their TSSs.

