## Supplementary Figures for "CHD4 conceals aberrant CTCF-binding sites at TAD interiors by regulating chromatin accessibility in mESCs"

Supplementary Figure 1

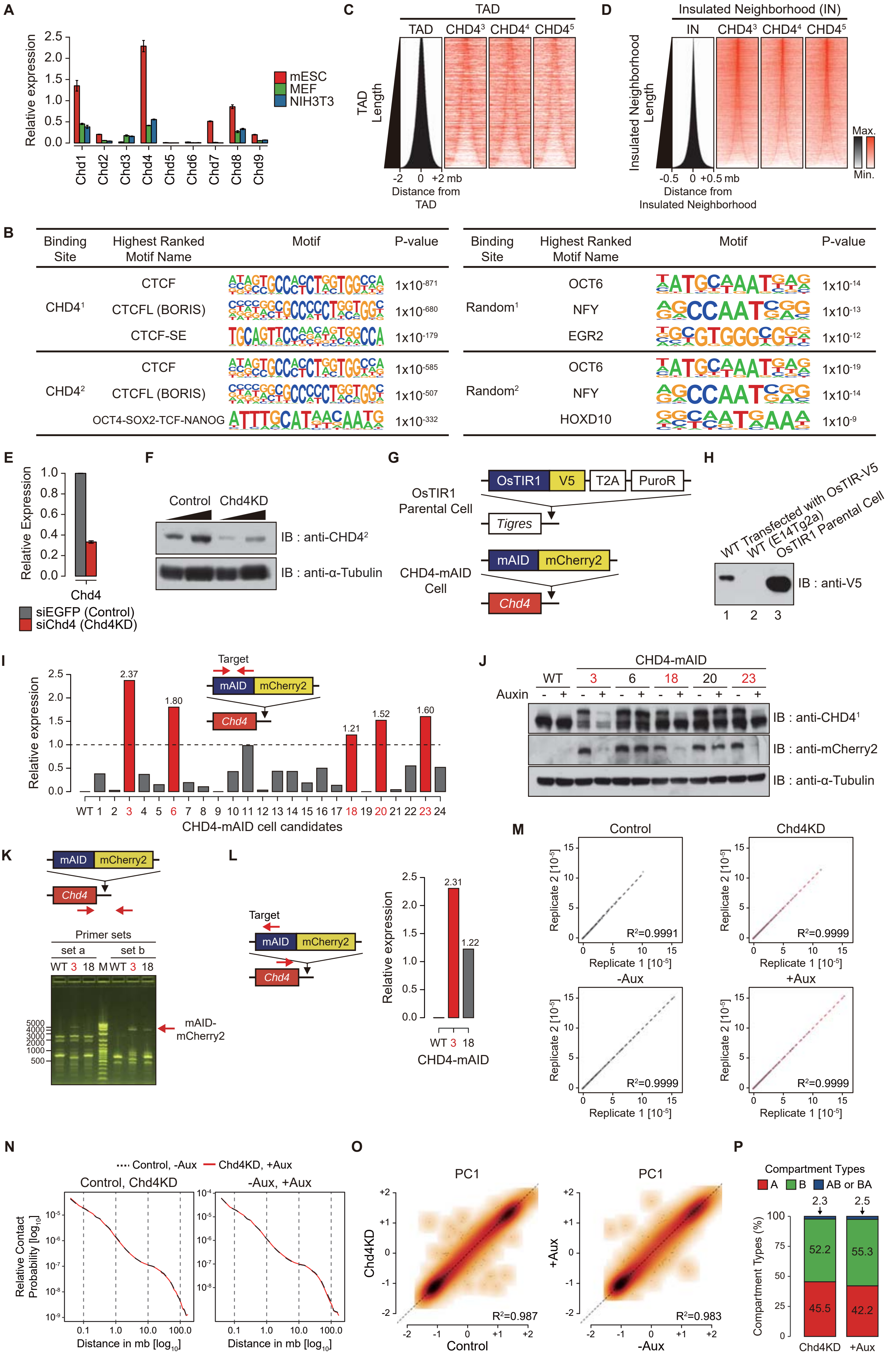

Supplementary Figure 2

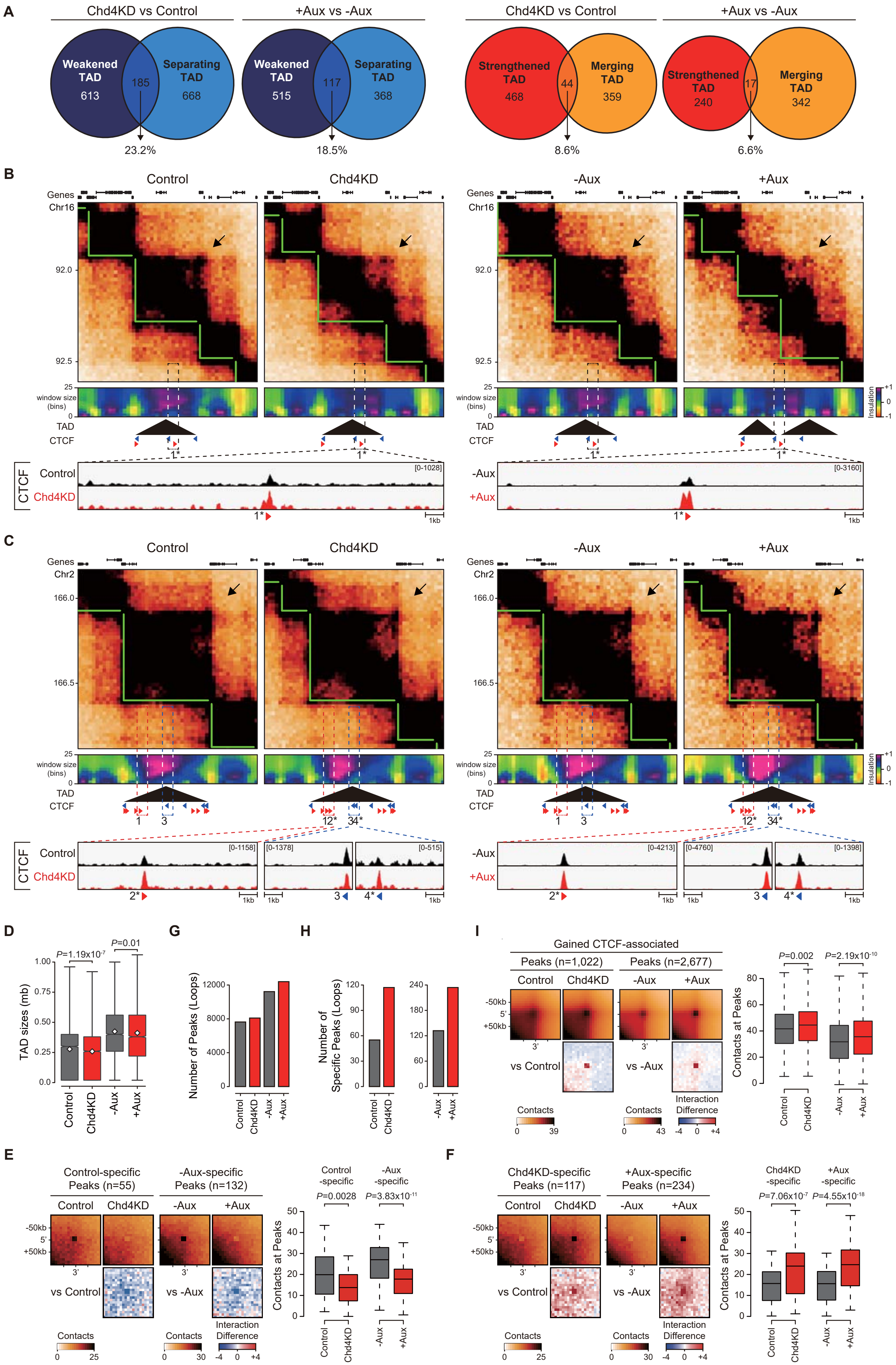

Supplementary Figure 3

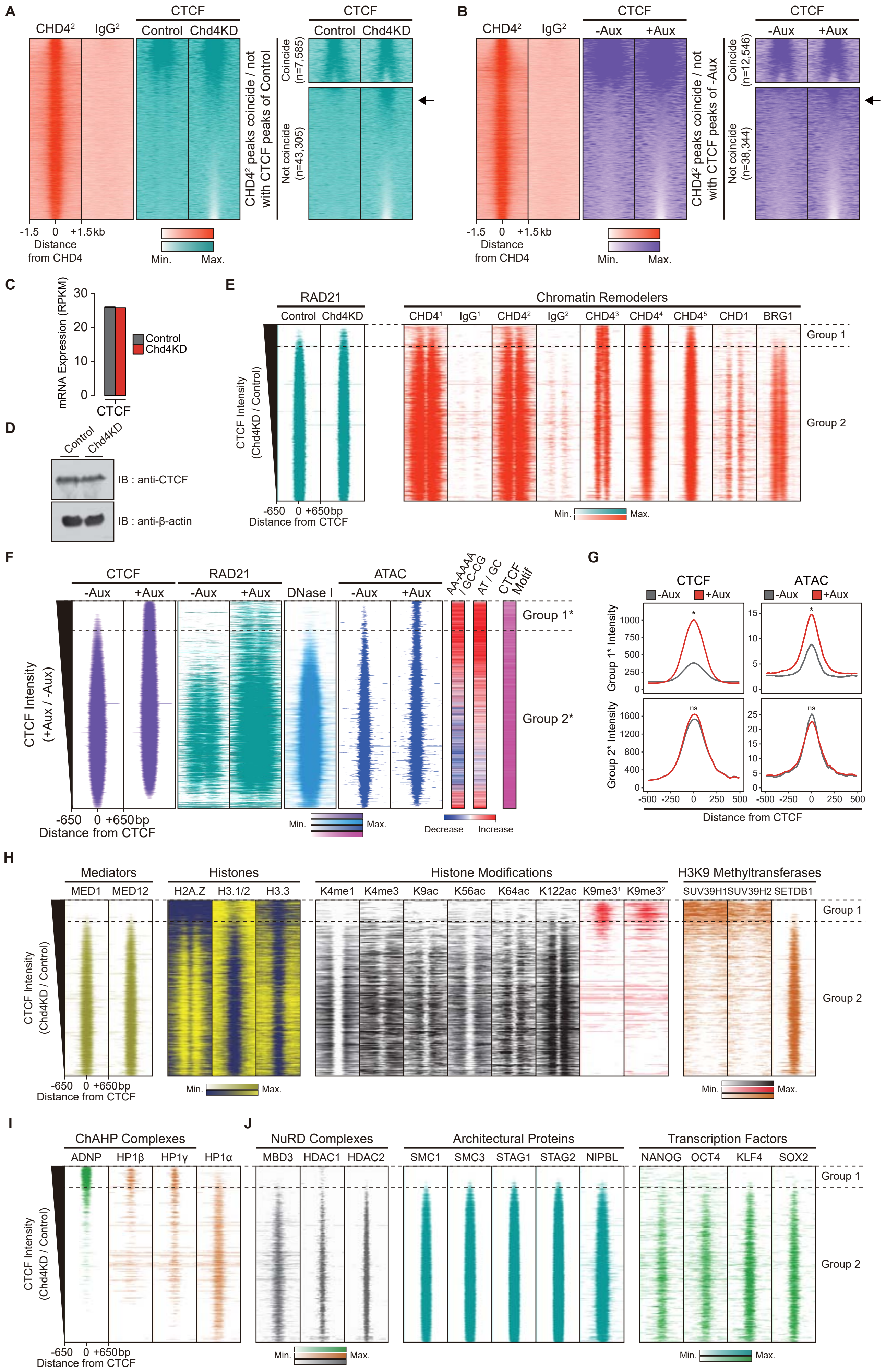

Supplementary Figure 4

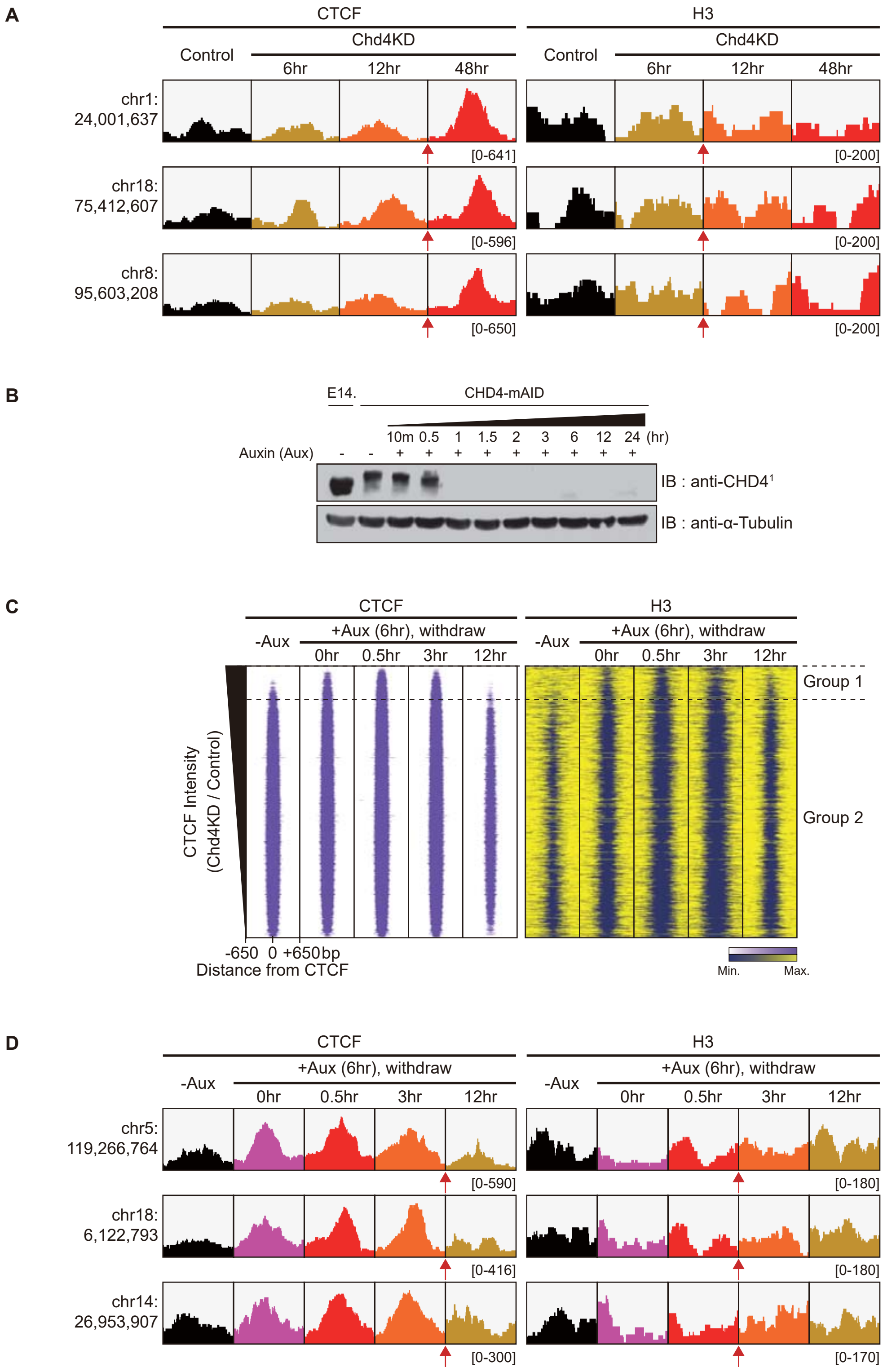

Supplementary Figure 5

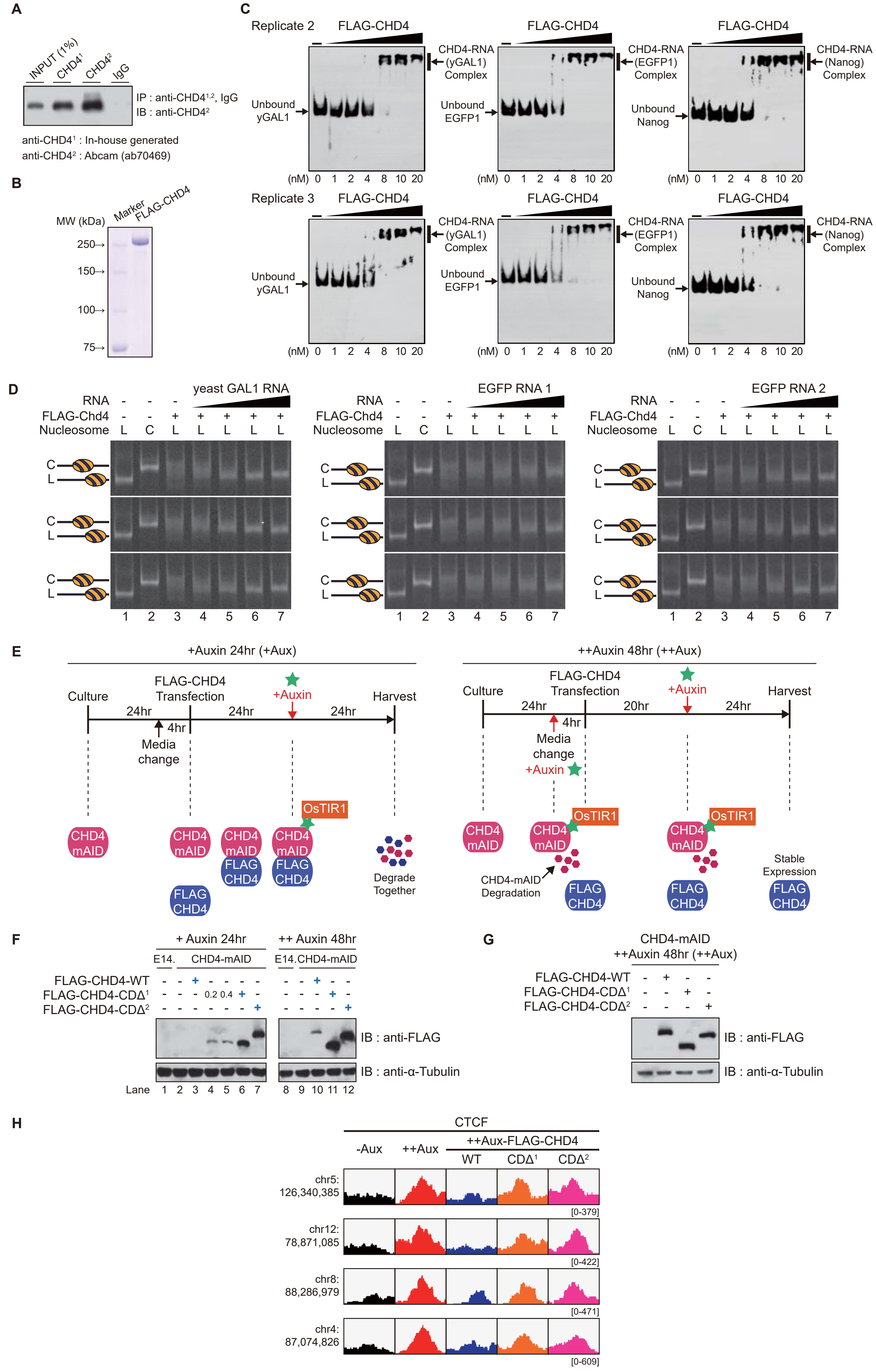

Supplementary Figure 6

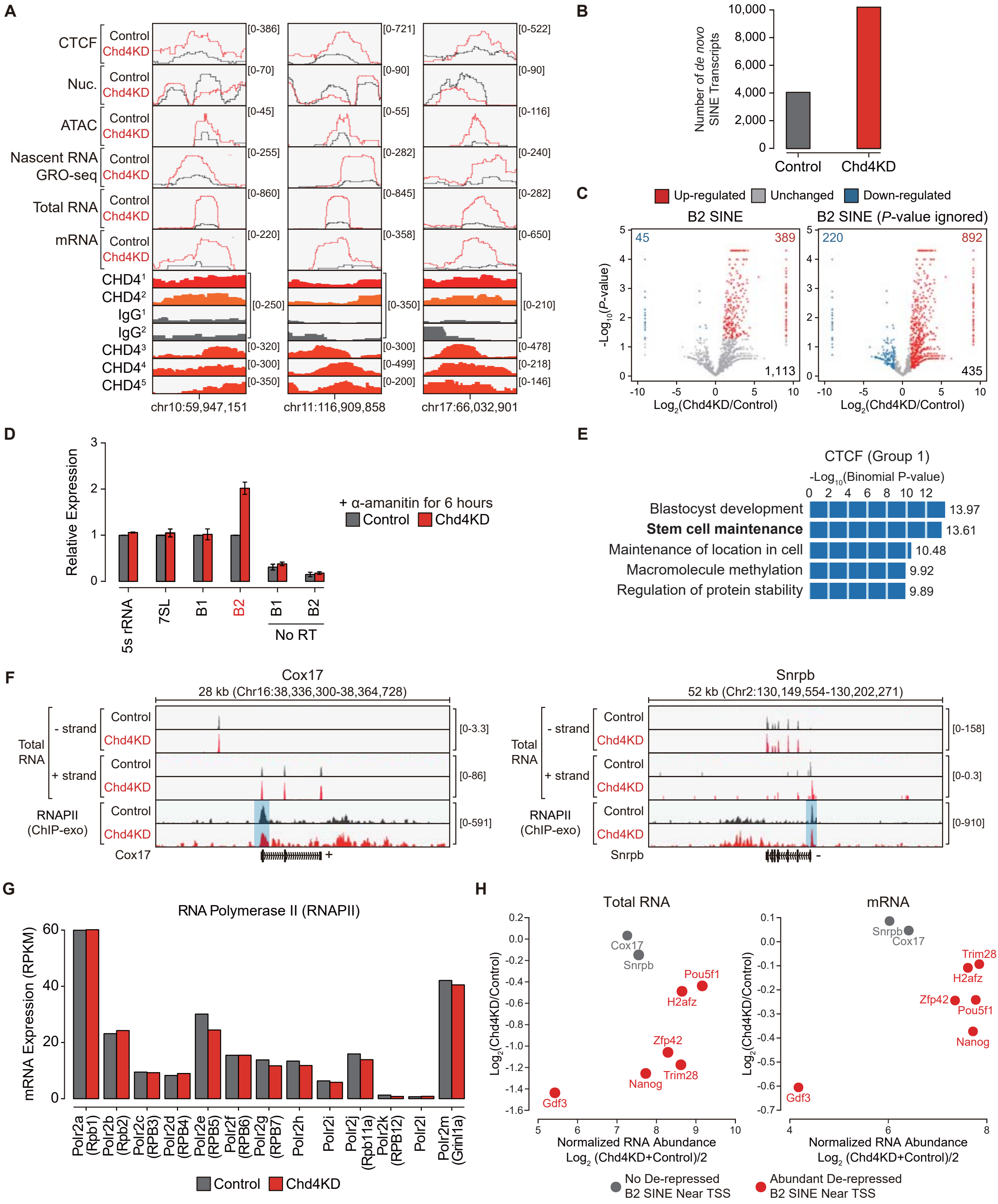
